# Stab Wound Injury Elicits Transit Amplifying Progenitor-like Phenotype in Parenchymal Astrocytes

**DOI:** 10.1101/2024.09.10.612217

**Authors:** Priya Maddhesiya, Finja Berger, Christina Koupourtidou, Alessandro Zambusi, Klara Tereza Novoselc, Judith Fischer-Sternjak, Tatiana Simon, Cora Olpe, Sebastian Jessberger, Magdalena Götz, Jovica Ninkovic

**Affiliations:** Biomedical Center Munich (BMC), Department of Cell Biology and Anatomy, Medical Faculty, LMU, Munich, Germany; Graduate School of Systemic Neurosciences, LMU, Munich, Germany; Research Unit Central Nervous System Regeneration, Helmholtz Center Munich, German Research Center for Environmental Health, Neuherberg, Germany; Institute of Stem Cell Research, Helmholtz Center Munich, German Research Center for Environmental Health, Neuherberg, Germany; Chair of Physiological Genomics, Biomedical Center (BMC), Faculty of Medicine, LMU Munich, Germany; Faculties of Medicine and Science, Laboratory of Neural Plasticity, Brain Research Institute, University of Zurich, Zurich, Switzerland; Munich Cluster for Systems Neurology SYNERGY, LMU, Munich, Germany

## Abstract

Astrocytes exhibit dual roles in central nervous system (CNS) recovery, offering both beneficial and detrimental effects. Following CNS injury, a subset of astrocytes undergoes proliferation, de-differentiation, and acquires self-renewal and neurosphere-forming capabilities *in vitro.* This subset of astrocytes represents a promising target for initiating brain repair processes and holds potential for neural recovery. However, studying these rare plastic astrocytes is challenging due to the absence of distinct markers. In our study, we characterized these astrocytic subpopulations using comparative single-cell transcriptome analysis. By leveraging the regenerative properties observed in radial glia of zebrafish, we identified and characterized injury-induced plastic astrocytes in mice. These injury-induced astrocytic subpopulations were predominantly proliferative and demonstrated the capacity for self-renewal and neurosphere formation, ultimately differentiating exclusively into astrocytes. Integration with scRNAseq data of the subependymal zone (SEZ) allowed us to trace the origins of these injury-induced plastic astrocytic subpopulations to parenchymal astrocytes. Our analysis revealed that a subset of these injury-induced astrocytes shares transcriptional similarities with endogenous transient amplifying progenitors (TAPs) within the SEZ, rather than with neural stem cells (NSCs). Notably, these injury-induced TAP-like cells exhibit distinct differentiation trajectories, favoring gliogenic over neurogenic differentiation. In summary, our study identifies a rare subset of injury-induced, proliferative plastic astrocytes with neurosphere-forming capacities. These cells originate from reactive astrocytes and resemble TAPs in their transcriptional profile. This study enhances our understanding of astrocyte plasticity post-injury.

**Highlights:**

- Single-cell transcriptomics and cross-species comparisons reveal proliferative and de-differentiated plastic astrocytes following CNS injury.
- Injury-induced de-differentiated astrocytes exhibit remarkable *in vitro* self-renewal and neurosphere formation but favor glial differentiation.
- De-differentiated astrocytes exhibit transcriptional similarities to transit-amplifying progenitors (TAPs) over neural stem cells (NSCs)
- Injury-induced TAP-like progenitors exhibit limited spontaneous neuronal differentiation.

## Introduction

Brain injuries, encompassing traumatic brain injuries (TBIs) and strokes, present formidable challenges to human health, resulting in enduring damage and functional deficits (Bramlett & Dietrich, 2015; Griesbach et al., 2018). These injuries highlight the limited regenerative capacity of the mammalian brain, which hampers the restoration of neural circuitry (Grade & Götz, 2017; Sun, 2014). Beyond disrupting functional neural circuits, brain injuries provoke intricate pathophysiological responses that culminate in the formation of a glial border (Sofroniew, 2009). This border acts as a physical barrier, isolating damaged tissue and preventing the spread of inflammation and further damage (Fawcett & Asher, 1999; Sofroniew, 2009). Astrocytes, microglia, and oligodendrocyte lineage cells undergo profound changes in morphology, gene expression, and function to establish and maintain the glial border (Liddelow & Barres, 2017; Matusova et al., 2023). Recent advancements have highlighted that certain component of the glial border, including subsets of reactive astrocytes, can promote axonal regeneration following spinal cord injury (Anderson et al., 2016). However, persistent neuroinflammation associated with the glial border alters the extracellular environment and impedes regeneration efforts (Y. Li et al., 2020; Sanchez-Gonzalez et al., 2022; Zambusi et al., 2022). A revolutionary approach seeks to transform glial cells proximal to injury into neurons, aiming to mitigate the detrimental effects of prolonged glial reactivity and provide new neurons for tissue repair in affected areas (Grade & Götz, 2017). Early *in vitro* studies have demonstrated the feasibility of converting glial cells into neurons with specific neurotransmitter identities through overexpression of neurogenic fate determinants (Berninger et al., 2007; Bocchi et al., 2022; Heinrich et al., 2010). Building on these findings, remarkable efficiency *in vivo* has been achieved in converting both astrocytes and NG2 cells into neurons (Liu et al., 2021; Mattugini et al., 2019; Pereira et al., 2017; Torper et al., 2015). Importantly, stab wound injury significantly enhances the conversion rate of parenchymal astrocytes into cells expressing neurogenic fate determinants compared to the intact brain (Mattugini et al., 2019). This aligns with recent findings indicating that subsets of astrocytes undergo identity shifts toward a more stem-like state following brain injury (Behrendt et al., 2012; Buffo et al., 2008; Dimou & Gotz, 2014; Gotz et al., 2015; Mori et al., 2005; Shimada et al., 2012; Simpson Ragdale et al., 2023; Sirko et al., 2023; Torper & Götz, 2017; Zamboni et al., 2020). Originally post-mitotic, these cells begin to proliferate and acquire the capacity to form multipotent neurospheres *in vitro* (Buffo et al., 2008; Gotz et al., 2015; Sirko et al., 2013). The mechanisms underlying astrocyte de-differentiation into neurosphere-forming cells after brain injury remain incompletely understood but several factors have been implicated to play a role. For instance, Sonic hedgehog (SHH) signaling induces stem cell responses in reactive astrocytes following invasive injuries, observed both *in vivo* and *in vitro* (Sirko et al., 2013). Notch signaling deficiency in cortical astrocytes enhances their neurogenic potential post-injury (Zamboni et al., 2020). Inhibition of Notch signaling increases the number and diversity of neurons derived from astrocytes in the striatum after stroke and improves motor function in mice, underscoring Notch’s role in maintaining the glial fate (Magnusson et al., 2014; Santopolo et al., 2020). Additionally, astrocytes bearing mutations in p53 generate more neurospheres than wild-type astrocytes after stab wound injury (Schmid et al., 2016; Simpson Ragdale et al., 2023), suggesting active mechanisms maintaining astrocyte identity. Sparse experimental evidence supports the hypothesis that injury induces transient de-differentiation of astrocytes with mechanisms preventing their neuronal differentiation. However, these lineage barriers can potentially be overcome by overexpressing neurogenic fate determinants following injury (Gascon et al., 2015, 2017; Heinrich et al., 2014; Mattugini et al., 2019), highlighting these astrocytes as promising targets for direct neuronal conversion. The prospect of utilizing plastic astrocytes as a source for generating new neurons raises significant concerns regarding their endogenous roles within the glial border. For example, proliferating astrocytes regulate monocyte trafficking post-injury, and interference with their function can lead to prolonged neuroinflammation (Frik et al., 2018). Astrocytes also play roles in blood-brain barrier recovery and neuroprotection following mild traumatic brain injury (George et al., 2022). Thus, identifying and understanding these cells’ lineage barriers and suitability as targets for direct conversion is crucial. However, prospective identification of plastic astrocytes has been challenging due to the lack of specific markers. Therefore, developing effective strategies to identify these rare injury-induced plastic astrocytes is essential for harnessing their potential in repairing brain injuries. In contrast to the mammalian brain, where astrocytes typically do not acquire neurogenic properties, the radial glia, astrocytic counterparts in zebrafish, exhibit plasticity and can differentiate into postmitotic neurons to facilitate endogenous repair after injury (Diotel et al., 2020; Zambusi & Ninkovic, 2020). Drawing from this observation, we hypothesized that plastic astrocytes in mice might share similarities with zebrafish radial glia. To investigate this hypothesis, we integrated single-cell RNAseq (sc RNAseq) data from intact and injured brains of zebrafish and mice. Our analysis revealed a subset of reactive astrocytes in mice that clustered closely with radial glia based on their transcriptomic profiles. These cells exhibited a unique transcriptional signature, characterized by high expression of the Ascl1 transcription factor. Using Ascl1 based genetic fate mapping, we demonstrated that these astrocyte-derived cells generate neurospheres following brain injury. Furthermore, employing pseudotime-based developmental trajectory analysis, we observed that these plastic cells transiently transition through a state resembling neural stem cells before adopting a gliogenic transit amplifying progenitor state. Our findings provide insights into the cellular and molecular mechanisms underlying the lack of endogenous neuronal generation in the injured mammalian brain. Specifically, we prospectively isolated plastic astrocyte-derived progenitors, characterized their distinct transcriptome, and identified lineage barriers that prevent their spontaneous differentiation into neurons. This study lays the groundwork for future investigations aimed at manipulating plastic astrocytes functionally to explore their native roles within the glial border and evaluate their potential for therapeutic brain repair strategies.

## Results

### Integration of single cell transcriptomes reveals shared cellular states in zebrafish and mouse brain

To identify rare injury-induced plastic astrocytic populations in the mammalian brain, we employed a trans-species approach. We hypothesized that murine plastic astrocytes would be similar to the zebrafish radial glia. Therefore, we integrated single-cell transcriptome data from both intact and injured mouse cerebral cortex and zebrafish telencephalon (Fig. 1A) (Koupourtidou et al., 2024; Zambusi et al., 2022). Specifically, we analyzed zebrafish brain cells at 3 and 7 days post-injury (dpi), corresponding to the onset (3 dpi) and peak (7 dpi) of radial glia reaction and proliferation (Baumgart et al., 2012; Sanchez-Gonzalez et al., 2022). Similarly, mouse cerebral cortex cells were isolated at 3 and 5 dpi, matching the onset of astrocytic proliferation (3 dpi) and the peak of neurosphere-forming capacity post-injury (5 dpi) (Buffo et al., 2008; Sirko et al., 2013). Using Seurat v4 (Butler et al., 2018) and a self-compiled function (see Methods), we integrated the zebrafish and mouse transcriptomes. Post-integration, cells from both species and conditions (injured or intact brain) were intermixed (Fig. 1B, Suppl. 1A). Unsupervised clustering with PCA (1:10) at a resolution of 0.7 revealed 25 distinct cell clusters (Fig. 1C). We annotated these clusters using cell type-specific markers, identifying various neuronal, macroglial, and microglial populations (Fig. 1C; Suppl. Table 1), including clusters expressing both astrocyte and radial glia (RG) markers, defined as Astrocyte/RG clusters (Fig. 1C). The distribution of mouse and zebrafish cells within these clusters indicated successful cross-species data integration (Fig. 1D-E).

**Figure 1.**
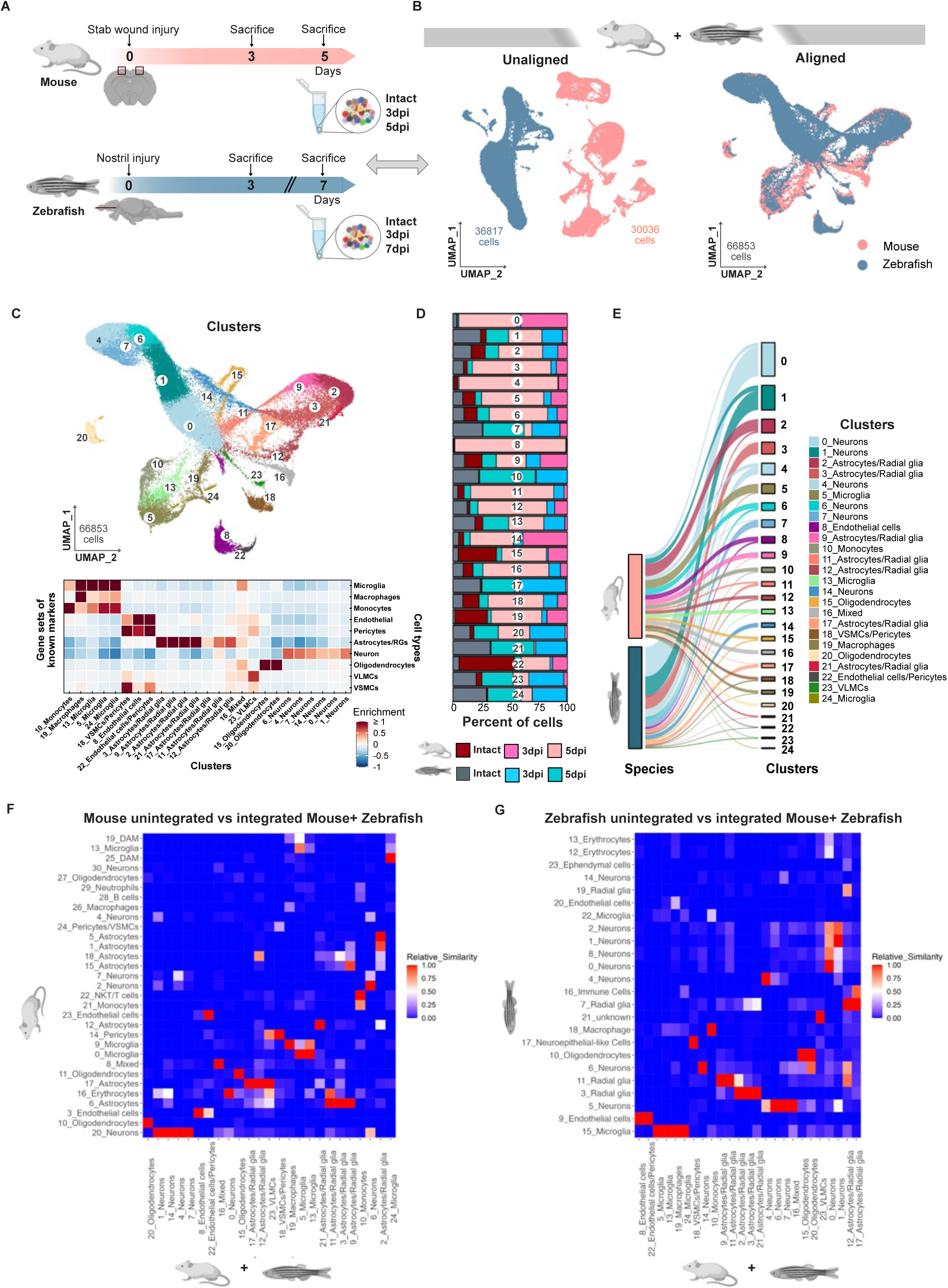
Integration of mouse and zebrafish single cell transcriptomes. (**A**) Schematic of datasets used for integration analysis from both species. (**B**) UMAP plots of unaligned (before integration) and aligned (after integration) datasets from a mouse (red) and zebrafish (blue). (**C**) UMAP plot depicting integrated dataset grouped into 25 transcriptionally distinct clusters, annotated by cell type-specific markers. (**D**) Bar plot depicting the contribution of cells from different conditions to identified cell cluster. (**E**) Alluvial plot visualizing contribution of the two species to identified clusters. (**F**,**G**) Heatmaps visualizing relative similarities among clusters between (F) unintegrated mouse (y-axis) and integrated mouse+zebrafish (x-axis) and (G) unintegrated zebrafish (y-axis) and integrated mouse+zebrafish (x-axis) datasets.

To validate our integration by independent method, we used the Harmony algorithm (Suppl. Fig. 1A-E), which relies on iterative integration and batch correction (Korsunsky et al., 2019). Harmony based unsupervised clustering identified 26 distinct cell clusters following PCA (1:10) at a resolution of 0.7 (Suppl. Fig. 1E). Importantly, this analysis showed similar results to Seurat based analysis, with most clusters containing cells from both species (Suppl. Fig 1B). We assessed cell type relationships between Harmony and Seurat clusters using ELeFHAnt’s *DeduceRelationship* function, evaluating relative cluster similarities (Thorner et al., 2021). Each Harmony cluster corresponded one-to-one with unique Seurat clusters (Suppl. Fig. 1C-E). Further analysis of the top 10 marker genes from Seurat cluster 2 Astro/RG and Harmony cluster 3 Astro/RG, which showed high relative similarity, revealed that 9 of the 10 top enriched genes were identical with similar enrichment (Suppl. Fig. 1F-I). These results suggest that intrinsic biological factors, rather than the integration algorithm, define the cell clusters in the integrated dataset. We extended our analysis by integrating intact and injured samples from both species, along with mouse peripheral blood mononuclear cells (PBMCs), using Seurat (Suppl. Fig. 2A). This allowed us to determine if the integration coerced distinct cell types into a unified representation. Our findings indicated that PBMCs clustered with brain immune cells (microglia and infiltrating monocytes), distinct from astrocyte/radial glia or neuronal clusters (Suppl. Fig. 2B-F; Suppl. Table 2), suggesting that the integration method did not enforce uniform clustering regardless of transcriptional features. To ensure that transcriptional information was retained during integration, we assessed the relative similarity between clusters in unintegrated mouse and zebrafish datasets and those in the integrated dataset using *DeduceRelationship* function of ELeFHAnt (Thorner et al., 2021) (Fig.1F, G). Our analysis showed a strong concordance between integrated dataset clusters and cell types in the unintegrated datasets, supporting the maintenance of specific cellular identities and essential transcriptional profiles. Notably, several clusters in the unintegrated datasets did not correspond to integrated clusters, suggesting the presence of species-specific cell types, which did not include astrocyte clusters and therefore did not affect our downstream analysis.

### A specific population of reactive astrocytes clusters with zebrafish radial glia

After integration, we were prompted to identify injury-induced plastic astrocytes as according to our hypothesis they would share the transcriptomic signature with zebrafish stem cells. Therefore, we focused our analysis on integrated clusters 2, 3, 9, 11, 12, 17, and 21 of Astrocytes/Radial glia (Astro/RG) and sub-clustered them further into a total of 10 Astro/RG sub-clusters using PCA (1:10) at 0.3 resolution (Fig. 2A, C). Indeed, the newly defined sub-clusters contained different proportions of cells originating from a specific condition (Fig. 2D, F). For example, cluster 0 was found to contain large proportion of cells originating from the intact mouse cerebral cortex (Fig. 2F). These cells also expressed the typical homeostatic astrocyte markers (Koupourtidou et al., 2024) in line with their origin (Fig. 2B, E, Suppl. Table 3). Interestingly, this cluster contained some zebrafish cells as well (Fig. 2D, F), suggesting that some radial glia could be more specialized to have a protoplasmic astrocyte function. Interestingly, cluster Astro/RG 0 also contained some astrocytes from the injured brains (Fig. 2B, F, E) in line with our previous observation that only a subset of astrocytes close to the injury shows reactive profiles (Koupourtidou et al., 2024). On the other hand, we identified Astro/RG 3 and 6 clusters which largely contained cells originating from zebrafish (Fig. 2D, F). The expression of typical proliferation genes (Fig. 2E) along with astroglial identity suggests that these cells belong to the actively cycling Type I radial glia (März et al., 2010). These clusters also contained cells from the injured mice but lacked cells originating from the intact mouse cerebral cortex (Fig. 2D, F). Importantly, a fraction of mouse Astro/RG cluster 3 and 6 cells also expressed the typical markers identifying this cluster as zebrafish Type I stem cells (März et al., 2010) (Fig. 2G), further highlighting the similarity of these mouse cells with the zebrafish stem cells.

**Figure 2.**
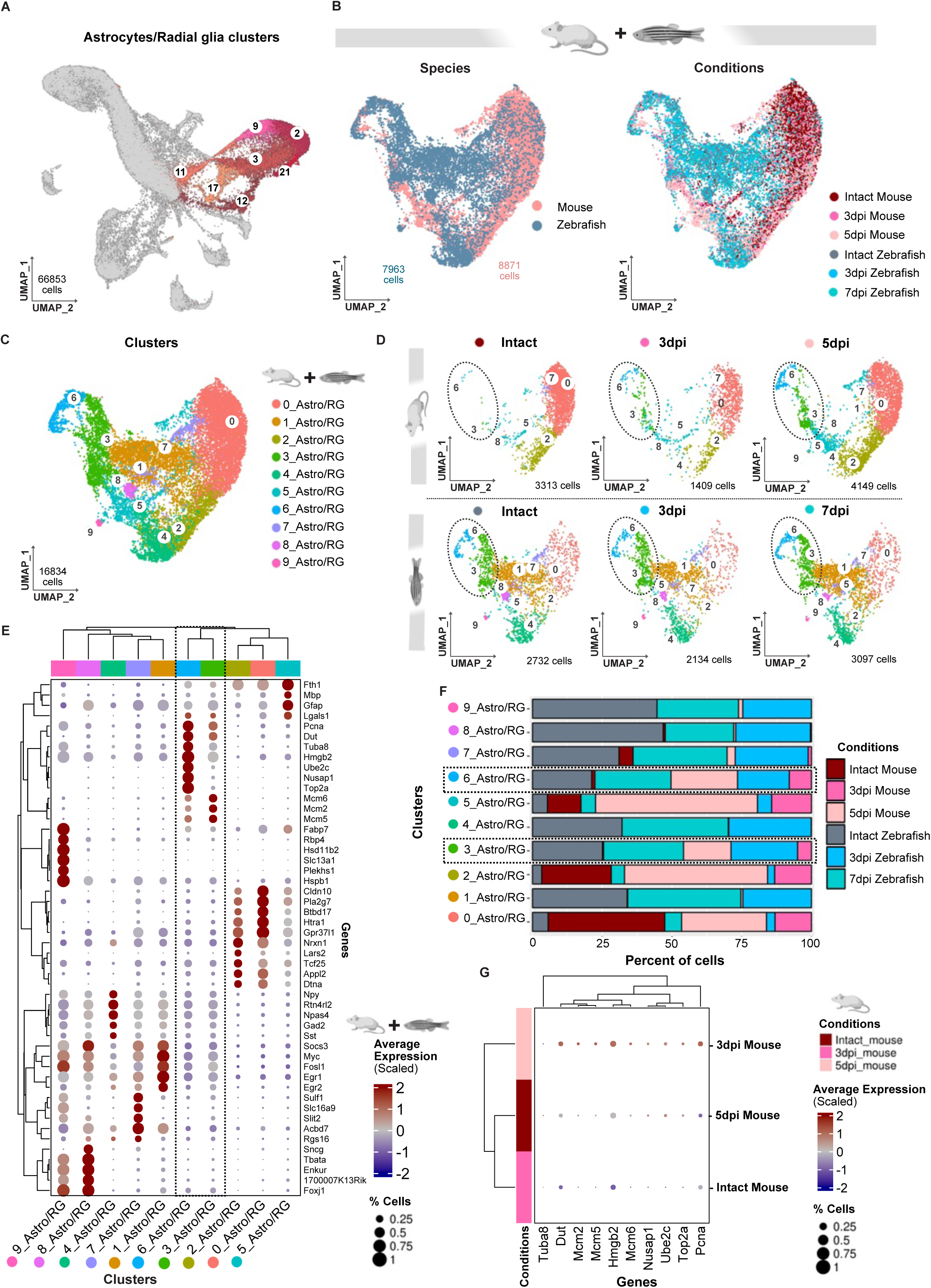
Integration of mouse and zebrafish species identifies Stab Wound injury-induced astrocytic population with radial glia properties. (**A**) UMAP plot highlighting astrocytes/radial glia (Astro/RG) in the integrated dataset. (**B**) UMAP plots visualizing cell distribution across species and conditions after sub-clustering of Astro/RG clusters. (**C**) UMAP plot depicting Astro/RG sub-clusters. (**D**) UMAP plots illustrating condition-specific distribution of cells within Astro/RG clusters mouse (above) and zebrafish (below). (**E**) Dot plot depicting top 5 enriched genes across Astro/RG sub-clusters. (**F**) Bar plot depicting cell distribution across injured and intact conditions in both species. Note that Astro/RG 3 and 6 clusters contain mouse cells only after injury. (**G**) Dot-plot showing the expression of top 5 enriched genes in Astro/RG 3 and 6 clusters, demonstrating elevated expression in injured mice (3 and 5 dpi) compared to intact mice. Note, only mouse cells from cluster Astro/RG 3 and 6 are considered for the analysis.

We further aimed at visualization of the Astro/RG 3 and 6 cluster cells in the injured tissue by examining the expression of genes enriched in these clusters. Our analysis revealed high expression of *Hmgb2*, *Uhrf1*, *Ascl1*, and *Rpa2* in the cluster Astro/RG 3/6 cells (Fig. 3A). These genes are upregulated in cells originating from the injured mouse cerebral cortex (both 3 and 5 dpi) compared to the intact mouse sample (Fig. 3B). Furthermore, we observed the increase in number of mouse cells expressing *Ascl1*, *Hmgb2* and *Uhrf1* at 5 dpi compared to 3 dpi (Fig. 3B, C). This increase in expression corresponds with the peak of astrocyte proliferation and neurosphere forming capacity (Sirko et al., 2013). The immunohistochemical and RNAScope analysis, showed that a subset of reactive astrocytes upregulates these genes in response to injury (Fig. 3 D-K; Suppl. Fig. 3 A-J) in line with observation in the scRNAseq that a proportion of cells expresses all of these genes at the single cell level (Fig. 3C). Importantly, a fraction of cells expressing HMGB2 also expressed *Uhrf1* (Fig. 3I) or *Ascl1* (Suppl. Fig. 3C). Notably, we also observed reactive, GFAP-positive astrocytes expressing only single marker genes (Hmgb2 or *Uhrf1*) (Fig.3 J,K and Suppl. Fig. 3J) in line with the hypothesis that Astro/RG 3/6 cluster cells upregulate the specific genes sequentially as they emerge from the homeostatic astrocyte state in response to brain injury.

**Figure 3.**
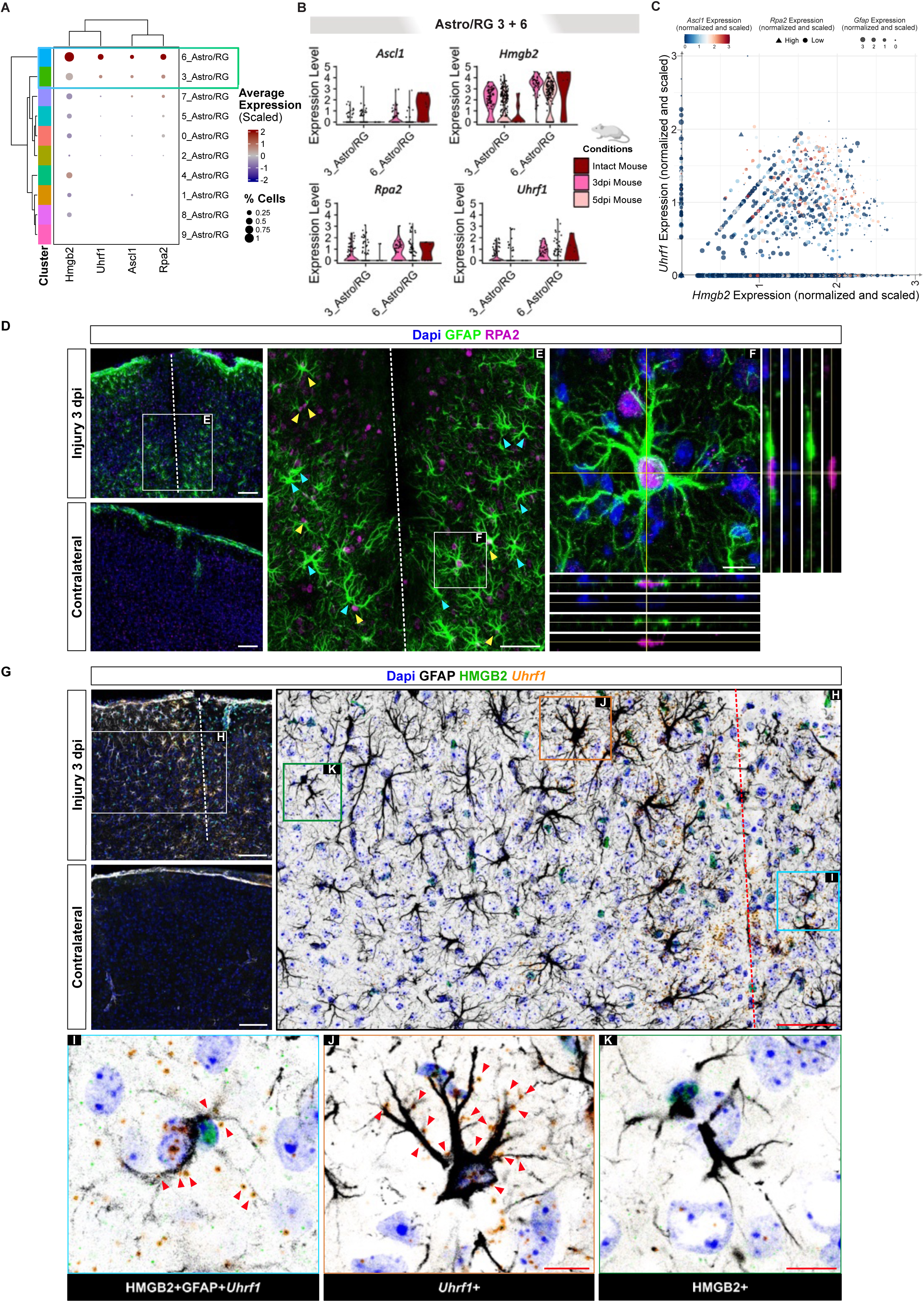
Visualizations of injury-induced Astro/RG clusters 3 and 6 cells after SW injury in mouse. (**A)** Dot plot showing the expression of Astro/RG 3/6 enriched genes in all Astro/RG sub-clusters in mice. (**B**) Violin plots illustrating the expression of indicated marker genes in sub-clusters Astro/RG 3/6 across conditions in mouse brain. (**C**) Plot illustrating the co-expression of Hmgb2, Uhrf1, Ascl1, Rpa2, and Gfap in mouse Astro/RG clusters. The X-axis and Y-axis represent normalized Hmgb2 and Uhrf1, respectively. Color indicates normalized Ascl1 expression (blue to red gradient), size represents normalized Gfap expression (larger points indicate higher expression), and shape differentiates Rpa2 expression (circles for ≤ 2, diamonds for > 2). (**D**) Micrographs depicting the astrocyte reactivity in the intact and injured (3 dpi) cerebral cortex based on the expression of GFAP, RPA2 and Dapi. (**E**, **F**) Expression of RPA2 in reactive astrocyte. Micrographs in E and F are magnifications of boxed areas in D and E, respectively. (**G-K**) Micrographs showing the co-localization of GFAP, HMGB2 and *Uhrf1* RNA in the intact and injured (3 dpi) mouse cerebral cortex. H is magnification of boxed area in G. Micrographs in I, J and K are single plane magnifications of boxed areas in H and depict a triple positive cell (I), an astrocyte (GFAP+) expressing only *Uhrf1* (J) and an astrocyte (GFAP+) expressing only HMGB2 (K). Red arrowheads in single optical sections (I-K) indicate *Uhrf1* molecules visible in the cell within that optical section. All other micrographs are maximum intensity projections of the confocal Z-stack and the micrograph in F contains orthogonal projections. Scale bars in D, G are 100 μm; 50 μm in E, H; 10 μm in F, I, J, K. White dashed lines (D, E, G) and red dashed line in H show position of the injury.

We next asked the question if the cluster Astro/RG 3/6 cells could be identified without the integration with zebrafish dataset (unintegrated analysis). Therefore, we clustered only astrocytes from the intact and injured mouse cerebral cortex and identified 7 distinct clusters at 0.3 resolution using PCA (1:15) (Suppl. Fig. 4A, B). We then identified cells from the cluster Astro/RG 3/6 in this unintegrated analysis. Indeed, we observed the distribution of the cluster Astro/RG 3/6 cells from the integrated analysis over 5 different clusters in the unintegrated analysis at resolution 0.5 (Suppl. Fig. 4E). Moreover, the different resolutions (0.3-0.8) of clustering also failed to fully isolate cluster Astro/RG 3/6 cells to the specific cluster in unintegrated analysis, despite clear enrichment in the case of cluster 6 (Suppl. Fig. 4 C-H), suggesting that this cellular state could only be isolated in integrative analysis.

### Cell proliferation is a hallmark of the injury-induced Astro/RG 3 and 6 clusters

The analysis of cluster-enriched genes in different astrocyte populations revealed a notable enrichment of cell proliferation-related genes within cluster Astro/RG 3/6 cells, including *Tuba8, Dut, Mcm2, Mcm5, Hmgb2, Mcm6, Nusap1, Ube2c, Top2a, Pcna* (Chen et al., 2021; G. Han et al., 2018; Kamino et al., 2011; Kimura et al., 2018; Nicolau-Neto et al., 2018; Ohtani et al., 1999; Ramos et al., 2020; Strzalka & Ziemienowicz, 2011; Wang et al., 2022; Xu et al., 2022; Yuan et al., 2022; Zeng et al., 2021) (Fig. 2E). Subsequent cell cycle analysis further revealed enrichment of distinct cell cycle phases across astrocytic sub-clusters (Fig. 4A). Notably, cluster Astro/RG 3 exhibited a significant proportion of cells in S phase (36.9% of all cluster Astro/RG 3 cells), while cluster Astro/RG 6 cells predominantly resided in G2M phase (93.3% of all cluster Astro/RG 6 cells) (Fig. 4B). Conversely, cells from homeostatic astrocyte clusters were largely in G1(G0) phase (Fig. 4A). Gene Ontology (GO) analysis underscored enrichment of processes associated with cell division, including translation, ribosome assembly, ribonuclear protein assembly, mitochondrial translation, and regulation of different phases of the cell cycle in both clusters Astro/RG 3 and 6 (Fig. 4C). Furthermore, examination of genes positively regulating the cell cycle (GO:0045787) revealed highest enrichment in Astro/RG clusters (Fig. 4D, Suppl. Table 4), suggesting a possibility that these two clusters contain de-differentiated cells in different phases of cell cycle. These finding suggests that cluster Astro/RG 3 and 6 contain astrocytes resuming proliferation in response to injury. To test this hypothesis, we labelled all cells undergoing cell division within the first 5 days after injury using BrdU incorporation (Fig. 4E). Reactive astrocytes were identified using GFAP immunoreactivity and cluster Astro/RG 3/6 astrocytes using their immunoreactivity for HMGB2 (Fig. 4F, G). Indeed, we observed that virtually all GFAP+ and HMGB2+ cluster Astro/RG 3/6 reactive astrocytes incorporated BrdU during the labelling period and only a few HMGB2+ and BrdU-cells were identified (Fig. 4G-J).

**Figure 4.**
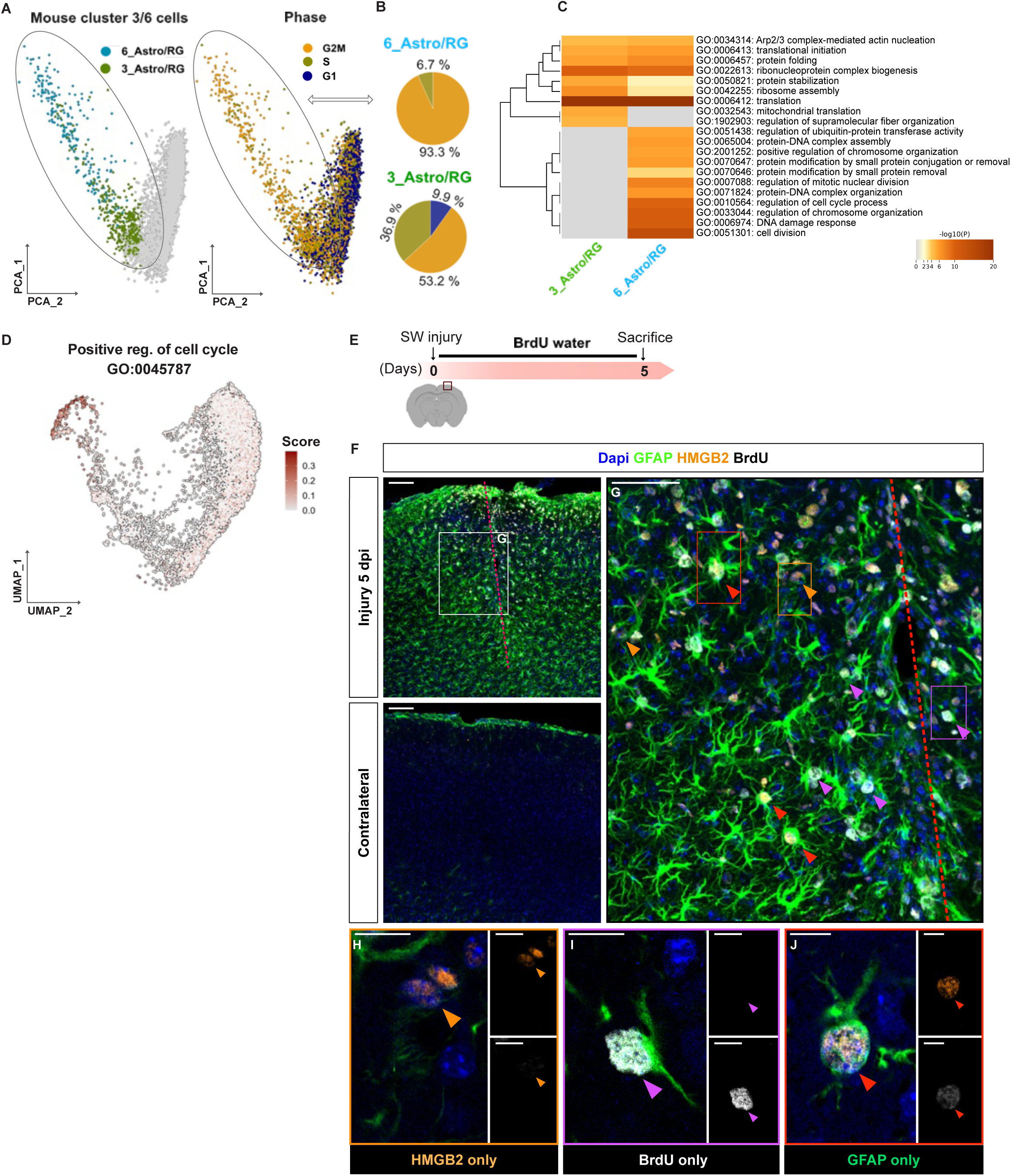
Astro/RG 3 and 6 clusters contain proliferative astrocytes. (**A**) PCA plots showing injury-induced mouse clusters (green and blue) and cell cycle phases (G1, S, G2M). (**B**) Pie charts depicting the distribution of Astro/RG 3 and 6 clusters amongst different cell cycle phases. (**C**) Heatmap showing GO terms enriched in injury-induced 3 and 6 Astro/RG clusters, colored by p-values. **(D)** UMAP plot displaying the enrichment score for genes associated with the Gene Ontology term “GO:0045787 positive regulation of cell cycle”. (**E**) A schematic illustrating the experimental design to address proliferation of Astro/RG 3 and 6 astrocytes using incorporation of BrdU. (**F**-**J**) Micrographs depicting BrdU incorporation by HMGB2+ reactive (GFAP+) astrocytes in the intact and injured (5 dpi) mouse cerebral cortex. G is magnification of boxed area in F. H-J are magnifications of boxed areas in G as indicated by color-code. Purple arrowheads indicate proliferating (BrdU+) reactive astrocytes (GFAP+), red arrowheads highlight proliferating (BrdU+) reactive astrocytes (GFAP+) also expressing HMGB2 and orange arrowheads point at reactive astrocytes (GFAP+) expressing only HMGB2. Micrographs in F and G are maximum intensity projections of confocal Z-stack. Micrographs in H-J are single optical sections. Scale bar in F is 100 μm; in G 50 μm and in H, I, J 10 μm. Red line indicates SW injury.

### Injury-induced cluster Astro/RG 3/6 astrocytes de-differentiate towards less mature state and generate neurospheres

As proliferative cluster Astro/RG 3/6 astrocytes emerged only after injury, we sought to understand their emergence by employing Monocle 3 (Cao et al., 2019). Monocle 3 enables the inference of temporal progression and cell fate decisions from scRNA-seq data. Pseudo-temporal ordering revealed the emergence of Astro/RG clusters 3 and 6 as continuum from the homeostatic astrocytes (Fig. 5A). The homeostatic astrocyte cluster 0 gives rise to the cluster Astro/RG 3/6 via intermediate clusters 2, 4 and 5. Interestingly, these clusters show the features of astrocyte reactivity, such as Gfap upregulation (Fig. 5B), but still do not have the proliferative features (Fig. 4D). The cluster Astro/RG 3 cells precede the cluster Astro/RG 6 cells (Fig. 5A), in line with cell cycle analysis, showing the larger fraction of cluster Astro/RG 3 cells being in S-phase and almost all cluster 6 cells undergoing G2M transition (Fig. 4A-B). This temporal analysis, therefore, suggests that the emergence of proliferative astrocytes after brain injury is a sequential continuum of transcriptional changes. This prompted us to analyze the expression of genes changing along the pseudo temporal trajectory. The typical astrocyte genes (*Aldoc, Gja1, S100b, Slc1a2, Slc1a3, Slc7a10*) decreased along the trajectory (Fig. 5B). *Gfap* first increased, reached the maximum in cluster 5 and then decreased as the trajectory approached clusters Astro/RG 3 and 6 (Fig. 5B). In contrast, we observed an increasing expression of genes associated with neural progenitors (*Ascl1, Dlx2, Olig2, Pcna, Hmgb2, Uhrf1*) (Fig. 5C). This data therefore suggests the gradual de-differentiation of protoplasmic astrocytes to reach the plastic, proliferative state. To test this hypothesis, we compared the transcriptomic profile of cluster Astro/RG 3/6 (proliferative clusters) and cluster 0 (homeostatic cluster, (Suppl. Fig. 5A)) to published transcriptomes of differentiated (AC1_RNA and AC2_RNA) and de-differentiated (TRP1_RNA and TRP2_RNA) astrocytes *in vitro* (Schmid et al., 2016). The gene set enriched in the cluster Astro/RG 3/6 astrocytes was also enriched in the de-differentiated TRP astrocytes, while the gene set identifying the homeostatic cluster Astro/RG 0 astrocytes shows enrichment in the homeostatic AC astrocytes (Suppl. Fig.5B). In addition, we compared the transcriptome of Astro/RG clusters with less mature cycling glial progenitors and astrocytes isolated from the postnatal (P4) mouse cerebral cortex (Di Bella et al., 2021). The similarity was assessed using the gene expression scores, defining cycling glial progenitors (cRGs cluster) and two astrocytic clusters (Astro_clust_1 and 2) in the P4 cortex (Suppl. Fig. 5C, Suppl. Table 5). Astrocytic clusters from the postnatal cortex shared similarities with homeostatic Astro/RG clusters, whereas Astro/RG clusters 3 and 6 exhibited resemblances to cycling glial cells (Suppl. Fig. 5A, C). These findings further substantiate the hypothesis that astrocytes undergo de-differentiation towards a less mature state (clusters Astro/RG 3 and 6) in response to injury.

**Fig 5:**
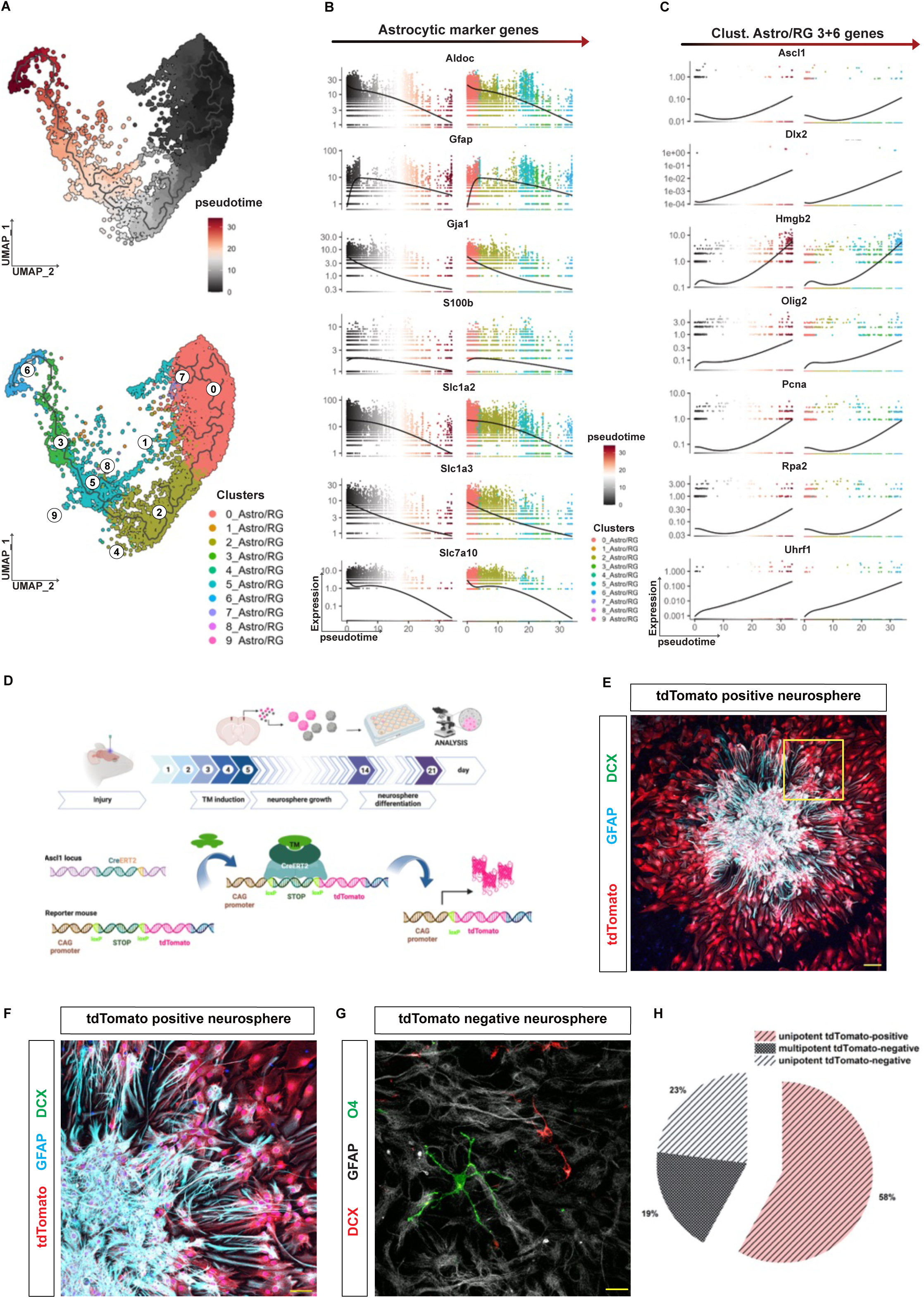
Injury-induced Astro/RG 3 and 6 cluster cells upregulate neural progenitor genes and gain neurosphere-forming potential. (**A**) UMAP depicting Monocle 3 pseudotime trajectories (upper plot) across Astro/RG clusters (lower plot) in the mouse. (**B**, **C**) Plots depicting the dynamic changes in the expression of astrocytic marker genes (B) and injury induced Astro/RG clusters 3 and 6 specific genes (C) along pseudotime trajectories. Note that Astro/RG clusters 3 and 6 specific genes are typical neural progenitor genes. (**D**) A schematic illustrating a neurosphere assay using Ascl1^CreERT2^ tdTomato mouse line. (**E-G**) Micrographs depicting reporter positive (E, F) and reporter negative (G) differentiated neurosphere stained for the lineage specific markers after 7 days in vitro. F is magnification of boxed area in E. All images are maximum intensity projections of the confocal Z-stack. Scale bars are 50 μm in E and 10 μm in F and G. (**H**) Pie chart depicting the differentiation potential of reporter positive and negative neurospheres (3 different animals have been analyzed).

As the immature neural progenitors and neural stem cells have the capacity to form neurospheres *in vitro*, we sought to test the capacity of cluster Astro/RG 3/6 cells to generate neurospheres. As Ascl1 marks these astrocytic clusters (Fig. 3A-C; Suppl. Fig. 3), we opted for Ascl1-based genetic fate mapping. Off note, Ascl1 is also expressed in oligodendrocyte progenitors (OPCs) in the mouse cerebral cortex regardless of brain injury. However, as PDGFRa positive OPCs do not form neurospheres (Buffo et al., 2008), we reasoned that any reporter positive neurospheres would be generated by Ascl1 positive cluster Astro/RG 3/6 astrocytes. For the genetic fate mapping we made use of a Ascl1^CreERT2^ mouse crossed to the tdTomato reporter mouse line, which expresses the red fluorescent protein tdTomato in Ascl1-expressing cells following tamoxifen treatment (Bottes et al., 2021; Madisen et al., 2010). The Cre-mediated recombination was induced 3 and 5 dpi based on the Ascl1 expression in pseudo temporal analysis (Fig. 3B, Fig. 5C) and cells were collected at 5 dpi for the neurosphere assay (Fig. 5D). As expected, we observed neurosphere formation only after brain injury. Importantly, about 60% of all generated neurospheres expressed the tdTomato reporter (Fig. 5E-H), suggesting that these neurospheres originated from the Ascl1-positive cluster Astro/RG 3/6 astrocytes. Interestingly, all reporter-positive neurospheres were unipotent and in the differentiation assay generated only astrocytes. In contrast, reporter negative neurospheres were both uni– and tri-potent in the differentiation assay (Fig. 5G, H). Taken together, we identified the injury-induced de-differentiated population of astrocytes with capacity to form unipotent neurospheres.

### Cluster Astro/RG 3/6 astrocytes display transcriptional features of several types of neural progenitors

The cluster Astro/RG 3/6 astrocytes appear to be unipotent in the neurosphere assay but still cluster with zebrafish neural stem cells possessing the capacity to generate neurons. Therefore, we reasoned that comparing their transcriptomes would identify the processes leading to unipotency. We identified 1123 (385 enriched in zebrafish and 738 enriched in mouse) differentially expressed genes (DEG) between zebrafish and mouse cluster 3 (Suppl. Fig. 5D, Suppl. Table 6) and 1089 DEGs (340 enriched in zebrafish and 749 enriched in mouse) in cluster 6 (Suppl. Fig. 5E, Suppl. Table 6). Collectively, zebrafish cells from Astro/RG 3 and 6 clusters exhibited enrichment in Wnt signaling, Notch signaling, G1 to S cell cycle control, ID-signaling and BMP signaling – all signaling pathways that have been implicated in regulation of neurogenesis in both zebrafish and mouse (Suppl. Fig. 5F, G). Conversely, cells from mouse Astro/RG 3 and 6 clusters showed enrichment in metabolic pathways, including oxidative stress, redox pathways, electron transport chain, glycolysis, and gluconeogenesis (Suppl. Fig. 5 F,G), suggesting that mouse cluster Astro/RG 3/6 astrocytes might fail to adopt their metabolic switch from astrocytes relaying on glycolysis to neural progenitors utilizing oxidative phosphorylation. As the specific metabolic programs appear to control the neuronal differentiation and neural stem cell maintenance in the adult mouse neurogenesis (Adusumilli et al., 2021; Beckervordersandforth et al., 2010; Wani et al., 2022), we hypothesized that incomplete transition of cluster Astro/RG 3/6 cells to neural stem cells might be the reason for the observed lack of potency and neurogenesis from cluster Astro/RG 3/6 astrocytes following injury. Therefore, we decided to compare the transcriptome of cluster Astro/RG 3/6 astrocytes and neural progenitors from the sub-ependymal zone in the adult mouse brain. We conducted an integrated analysis by combining single-cell transcriptome data from the SEZ of adult mice with previously collected data from both intact and injured (3 and 5 dpi) cerebral cortex (Fig. 6A). This approach allowed us to identify major cell types, including astrocytes, oligodendrocytes, microglia, transient amplifying progenitors (TAPs), neuroblasts (NBs), and neurons (Suppl. Table 7, Fig. 6B, D). However, quiescent adult NSCs share many markers with astrocytes, making it difficult to delineate these two cell types in the integrated analysis (Fig. 6C, D). Therefore, we performed a separate analysis focusing exclusively on the SEZ condition (Suppl. Fig. 6A). Within this analysis, we identified distinct populations, including quiescent NSCs (qNSCs), activated NSCs (aNSCs), TAPs, and NBs (Suppl. Fig. 6B-D) based on known markers (Suppl. Table 7). Additionally, we observed continuous pseudotime trajectories from quiescent NSCs to NBs, reflecting the inherent differentiation process of NSCs (Suppl. Fig. 6E). Furthermore, when we mapped SEZ NSC cells (qNSCs and aNSCs) back to the integrated and subclustered astrocyte clusters alongside TAPs and NBs (Suppl. Fig. 6F), we confirmed the presence of quiescent NSCs and activated NSCs within the integrated astrocyte clusters, validating their coexistence and affirming the robustness of our analysis.

**Figure 6.**
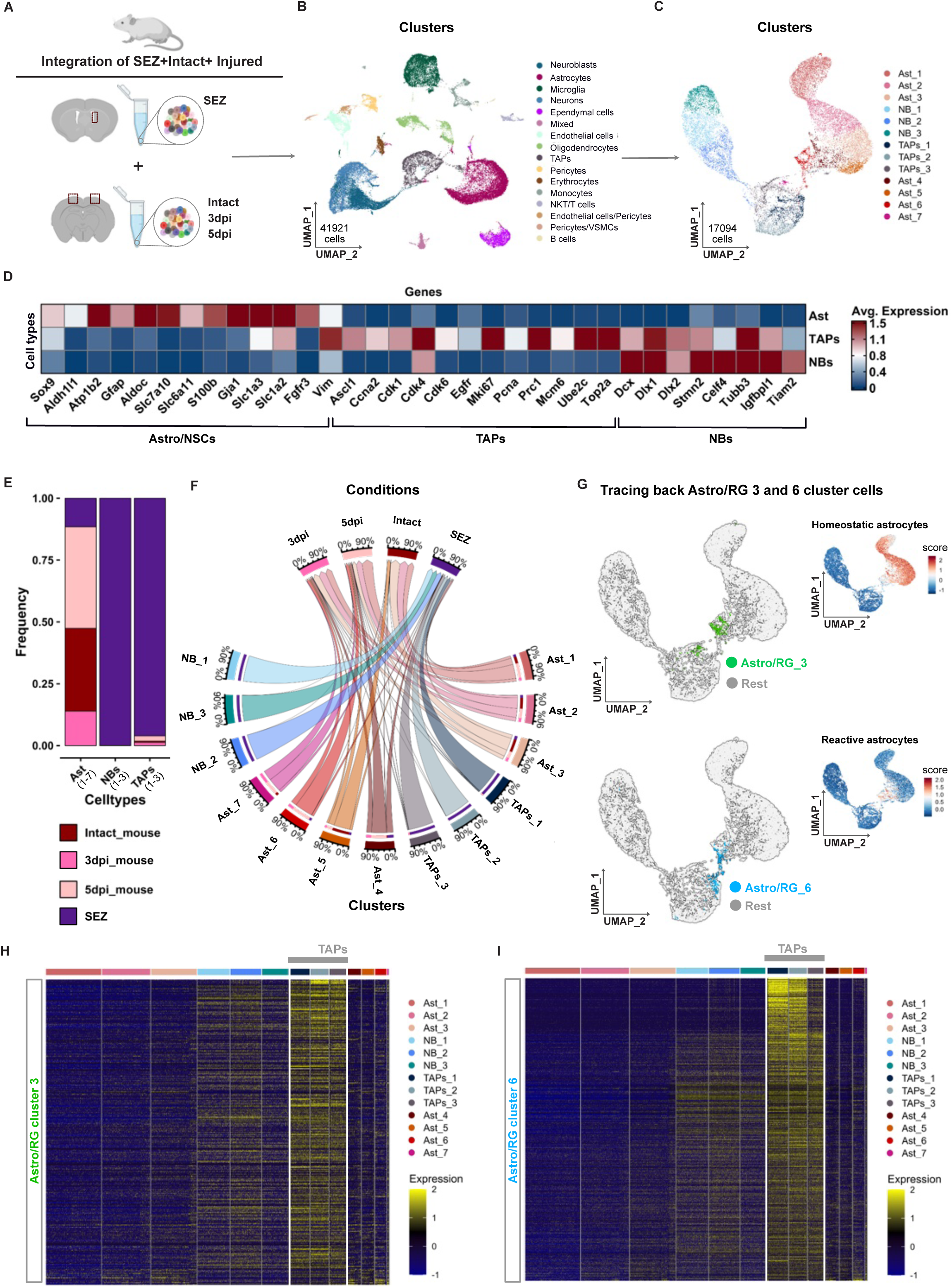
Some injury-induced Astro/RG 3 and 6 cluster cells show similarities to Transit Amplifying Progenitors (TAPs). (**A**) schematic illustrating datasets used for the integration analysis. (**B**) UMAP plot displays single cells grouped into 16 distinct cellular clusters annotated using known cell-type-specific markers. (**C**) UMAP plot demonstrating sub-clustering of the neurogenic lineage cells containing astrocyte/neural stem cells (Ast), TAPs and neuroblast (NBs) clusters. (**D**) Heatmap depicting expression of known cell type markers across Ast, TAPs, and NBs (Suppl. Table 7). (**E**) Bar plot illustrating frequency distribution of cells from cortex and SEZ conditions across Ast, TAPs, and NBs cell types. (**F**) Circos plot showing the distribution of cells from different conditions amongst specific Ast, TAPs, and NBs clusters. (**G)** UMAP plot showing the cells of injury-induced Astro/RG 3 and 6 clusters (from the zebrafish/mouse integration) identified in the integrated cortex and SEZ dataset. Inlets represent the enrichment score for homeostatic and reactive astrocytes calculated based on the gene expression published by Koupourtidou et al. 2024. (**H**, **I**) Heatmaps representing similarities of the transcriptome of injury-induced cluster Astro/RG_3 (H) and Astro/RG_6 (I) with SEZ Ast, TAPs, and NBs.

Moreover, this allowed us to assess the congruence among de-differentiated Astro/RG 3 and 6 cluster cells, TAPs, and NBs within the integrated SEZ+cortex analysis. In line with the absence of restorative neurogenesis in the cortex following injury (Buffo et al., 2008), we did not observe any cells from the cerebral cortex in the cluster containing SEZ neuroblasts, while clusters containing stem cells and TAPs contained cells from SEZ, intact and injured cortex (Fig. 6E-F). Furthermore, cross-referencing identities confirmed presence of cluster Astro/RG 3/6 cells in several clusters with astrocyte identity Ast_4, Ast_6 and Ast_7 (Fig. 6G). To our surprise, we observed that 20 % of cluster 6 cells clustered with SEZ derived TAPs_1 (Fig. 6F, G). Importantly, the TAPs_1 cluster did not contain any cells from the intact cerebral cortex (Fig. 6F). This prompted us to compare the transcriptome of the cluster Astro/RG 3/6 cells and neurogenic lineage cells identified in the SEZ only analysis following cell cycle gene regression. Indeed, we observed that both cluster Astro/RG 3 and 6 cells show the highest transcriptional similarity to the clusters of TAPs (Fig. 6H, I). Taken together, our analysis suggests that the injury-induced, de-differentiated cluster Astro/RG 3/6 astrocytes spread along the neurogenic lineage acquiring features of several progenitor types.

### Injury-induced, plastic astrocytes differentiate to TAP-like state

The distribution of cluster Astro/RG 3/6 cells along the neurogenic lineage prompted us to delineate their differentiation path using pseudotime trajectory and diffusion map analyses (Fig. 7A and Suppl. Fig. 7A, B). To differentiate between cortex and SEZ cells in the integrated object, the pseudotime was performed within the integrated object but considering either only SEZ or only cortical cells (Fig. 7A-C). As expected, the differentiation trajectory for SEZ cells started at the cluster containing qNSCs (Ast_1 cluster), went via aNSCs-containing cluster to TAP-containing clusters and ended up in the neuroblasts-containing cluster (Fig. 7B). Interestingly, we did not observe the heterogeneity represented by different clusters only in NSCs, but also in the TAP population. The TAPs_3 cluster transitioned into NBs, while TAPs_2 and TAPs_1 showed higher enrichment for proliferation markers (Figure 7B, Suppl. Fig. 7F). In the cortex, pseudotime trajectory analysis unveiled a shift from homeostatic astrocytes to reactive astrocytes and subsequently to the TAPs_1 cluster (Fig. 7C). Remarkably, based on trajectory analysis, TAPs_1 cortical cells were not found to contribute to the trajectory of neuroblasts clusters (Fig. 7A-C). To confirm these state transitions by an independent method, we performed diffusion map analysis (Suppl. Fig. 7A, B). In the SEZ, we found three distinct states corresponding to NSCs (q/a), TAPs and NBs with transitions identical to the pseudotime analysis (Suppl. Fig. 7A). In the cortex, we also identified three clusters of cells corresponding to homeostatic astrocytes, reactive astrocytes, and TAPs (Suppl. Fig. 7B), further supporting an emergence of TAP-like state from the homeostatic astrocytes via reactive astrocyte clusters that is similar but not identical to aNSCs following brain injury.

**Fig 7:**
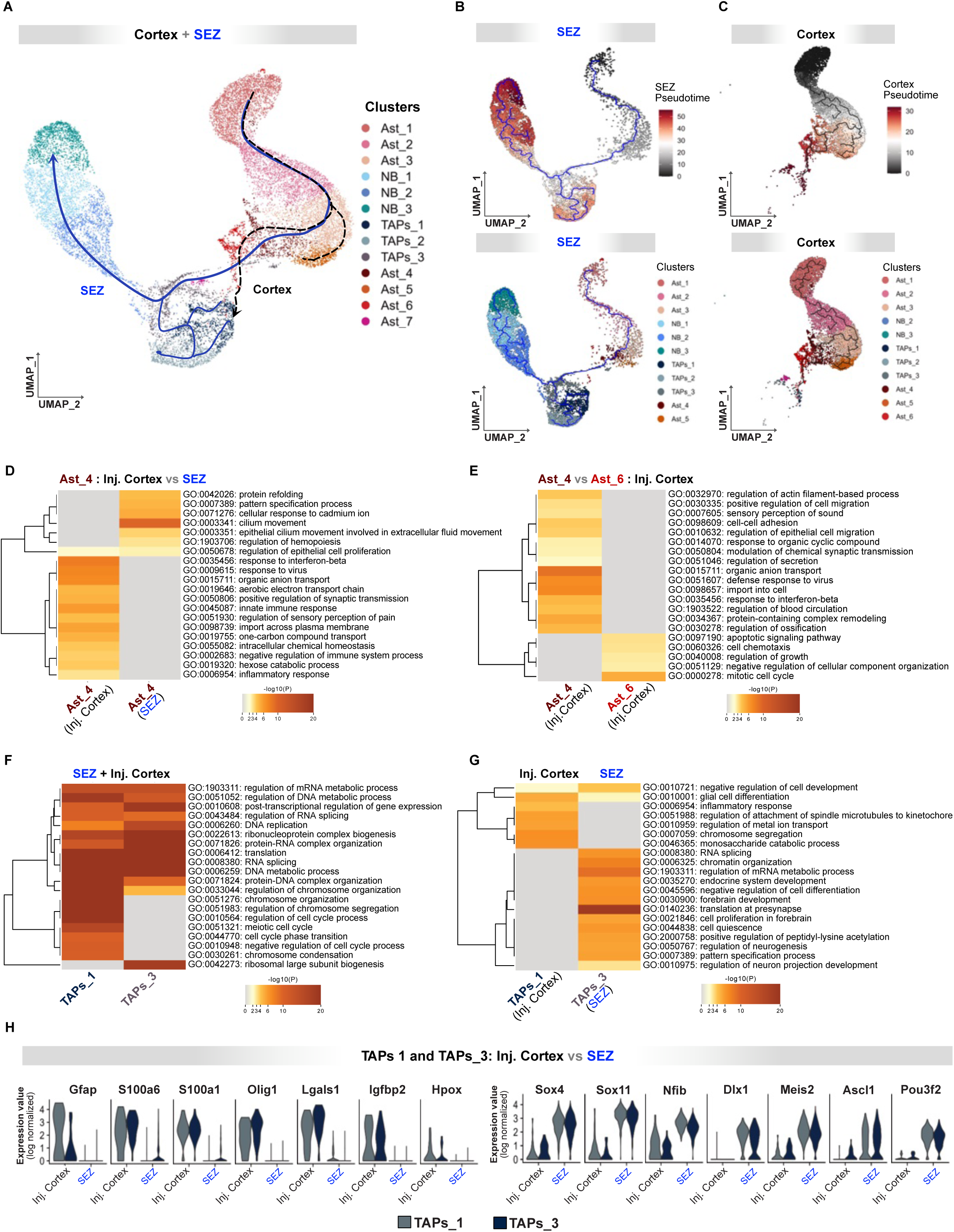
TAP-like cells emerging after injury fail to upregulate neurogenic fate determinants. (**A**) UMAP plot illustrating representative pseudotime trajectory in the SEZ (solid blue line) and injured cerebral cortex (dotted black line) (**B**, **C**) UMAP plots of pseudotime trajectory of SEZ only cells (B) and cerebral cortex only (C) based on pseudotime (upper panels) and across clusters (lower panels). (**D**) Heatmap depicting enriched GO terms in the set of DEGs between Ast_4 cluster from the injured cerebral cortex and SEZ, color-coded by p-values. (**E**) Heatmap illustrating enriched GO terms in the set of DEGs between Ast_4 and Ast_6 clusters isolated from the injured cerebral cortex, color-coded by p-values. (**F**) Heatmap of enriched GO terms in the DEG set between TAPs_1 and TAPs_3 clusters of the integrated cortex and SEZ, colored by p-values. (**G**) Heatmap of GO terms enriched in DEGs between TAPs_1 of the injured cortex and TAPs_3 of SEZ, colored by p-values. (**H**) Violin plot displaying 7 significant DEGs between injured context and SEZ in TAPs_1 and TAPs_2 clusters, color-coded by TAPs clusters.

The SEZ and cortical trajectories diverged at the level of astrocytic cluster Ast_4 (Fig. 7A). DEG analysis of SEZ and cortical cells contributing to Ast_4 showed an enrichment of GO terms related to cilium movement, pattern specification processes, epithelial cilium movement, and protein refolding in the SEZ cells (Fig. 7D). These processes are known to be associated with stem cell differentiation and renewal (Moore et al., 2015; Yanardag & Pugacheva, 2021). Conversely, Ast_4 cells from the injured cortex exhibited enrichment in GO terms such as inflammatory response, response to virus, innate immune response, and interferon beta response (Fig. 7D). As these are the terms linked to the astrocyte reactivity (Koupourtidou et al., 2024), this suggests that the cells from the injured cortex did not completely downregulate inflammatory, injury-induced programs and failed to establish neural stem cell maintenance network. This is in line with the observation that injury-induced astrocyte plasticity diminishes after 7 days (Buffo et al., 2005, 2008). Moreover, the SEZ trajectory transits from the Ast_4 directly to the TAP clusters, while the cortical trajectory contains one additional astrocytic cluster, the cluster Ast_6 (Fig. 7A-C). The direct comparison of cortical cells from the Ast_4 and Ast_6 clusters revealed an enrichment of GO terms related to inflammatory response (interferon-beta response, defense response to virus) in the Ast_4 cells, while processes associated with proliferation were enriched in the Ast_6 cluster (Fig. 7E). Together, these findings suggest that the additional astrocytic state detected in the cortical trajectory could be due to longer time that these cells need to downregulate the inflammatory processes. Once the inflammatory processes are downregulated, they could proceed further to the TAP state (TAPs_1) and upregulate processes necessary for the proliferation.

### Injury-induced TAPs fail to upregulate neurogenic fate determinants

The progression towards TAP states was associated with the expression of typical TAP markers such as *Ascl1, Dcx, Olig1, Olig2*, and *Mki67* (Suppl. Fig.7F, G) in both pseudotime trajectories. Additionally, we observed a decline in the expression of astrocytic markers (e.g., *Sox9* and *Slc1a2*; Suppl. Fig.7 F, G) within these clusters as they transited into TAP-like state. The GO term analysis revealed that TAP clusters (TAPs_1 and TAPs_3) activate processes linked to metabolism, replication, post-translational gene expression regulation, and translation (Fig. 7F), in line with the reported need for metabolic changes and translation regulation along the neurogenic lineage (Adusumilli et al., 2021; Baser et al., 2017, 2019; Beckervordersandforth et al., 2017; Knobloch et al., 2017; Wani et al., 2022). However, the cortical cells from the TAPs cluster were not observed to continue along the neurogenic lineage towards neuroblasts (Fig. 7C). Therefore, we conducted DEG analysis between the neurogenic TAPs_3 from the SEZ and cortical TAPs_1 cells (TAPs without the transition to neuroblasts) (Fig.7G, Suppl. Table 8). Our analysis revealed that injury-induced TAP-like clusters still expressed glial-associated genes (e.g., *Gfap, S100a1, S100a6, Olig1, Lgals1, Igfbp2*) as well as NSCs markers (*HopX*)(D. Li et al., 2015) (Fig. 7H), implicating that they failed to completely erase their previous states. Moreover, we observed that they did not upregulate typical neurogenic genes (*Sox4, Sox11, Nfib, Dlx1, Meis2, Pou3f2*) that are however upregulated in the SEZ TAP trajectory (Fig. 7H). Moreover, the cortical TAPs_1/TAPs_2 cluster cells express high levels of genes indicative of Notch pathway activation (Suppl. Fig. 7C-E, Suppl. Table 9). Importantly, these levels are comparable with the Notch activity levels in the bona fide neural stem cell clusters (Suppl. Fig. 7D, E). This is line with findings that Notch activity inhibits progression of neural stem cells towards neurogenic progenitors (Imayoshi et al., 2010) and reports that inhibition of Notch in the astrocyte-derived cells after brain injury allows their differentiation to neurons (Zamboni et al., 2020). The analysis of expression of specific lineage genes was further confirmed by the unbiased GO term analysis. Genes specifically expressed in the TAPs_3 cluster were enriched in the processes related to neurogenesis, while genes specifically enriched in the TAPs_1 cluster were related to inflammatory response, monosaccharide catabolic process, chromosome segregation, and metal ion transport (Fig. 7G). Taken together, our analysis proposes that injury induces the de-differentiation of post-mitotic astrocytes towards a state similar to aNSC-like state. However, these cells fail to generate properly specified TAPs lacking the expression of critical neurogenic genes and, therefore, hindering further lineage progression towards neuroblasts.

## Discussion

### Multi-species data integration

Cell lineage barriers largely define the cellular reaction to different brain pathologies, including stab wound injury (Gascon et al., 2017; Ninkovic & Götz, 2018). Pathology-induced crunching of these cellular barriers is the basis for the glial cell reactivity following brain pathology as well as their experimental trans-differentiation for repair purposes. Importantly, glial cells show different levels of barrier plasticity, with astrocytes showing the most drastic changes. Namely, a subset of originally post-mitotic astrocytes re-enters the cell cycle, expresses NSCs markers, gains the capacity to self-renew and generates multipotent neurospheres *in vitro* (Sirko et al., 2013). Such a dramatic change in the cell and molecular biology of astrocytes in response to insult brings an important question about the functional importance of this astrocytic population. The main caveat in addressing this question is the prospective isolation of these cells. Indeed, several studies identified the plastic, proliferative astrocytes retrospectively in both, animal model organisms (Bardehle et al., 2013; Lange Canhos et al., 2021; Sirko et al., 2015) and postmortem human brain (Sirko et al., 2023), making it difficult to specifically modify their reaction after injury and address their function. The recent advances in single cell profiling technologies did not really resolve this problem despite the identification of enormous astrocyte heterogeneity in both healthy and pathological conditions (Batiuk et al., 2020; Bugiani et al., 2022; Clarke et al., 2018; D’Elia et al., 2023; R. T. Han et al., 2021; Holt, 2023; Liddelow et al., 2017; Matias et al., 2019; Schober et al., 2022), even including the identification of proliferative astrocytes with stem cell characteristics in the intact diencephalon (Ohlig et al., 2021). One possible explanation for this could be that the currently available methods to prepare the single cell suspension specifically miss this astrocytic population, as the retrospective characterization of de-differentiated, proliferative astrocytes revealed the particular localization of these cells to the juxtavascular compartment (Bardehle et al., 2013). In addition, the proliferative astrocytes are a small cellular population that could be missed due to the lack of power of currently available datasets (R. T. Han et al., 2021). To overcome these limitations, we have recently optimized a cell isolation method for single cell transcriptome analysis (Koupourtidou et al., 2024) that recovers most of the glial cells and reveals more glial heterogeneity compared to so far available datasets (Koupourtidou et al., 2024). Moreover, we paired this analysis with the trans-species data integration to increase the power of our analysis. Indeed, this approach led to the identification of the specific cluster composed largely of zebrafish radial glia with stem cell properties. In addition, this cluster contained a small fraction of astrocytes from the injured tissue in line with the hypothesis that the de-differentiated astrocytes in the cerebral cortex could be observed only after brain injury (Sirko et al., 2013). Importantly, a separate cellular cluster of proliferating plastic astrocytes could not be identified using only the dataset from the mouse brain as the cells were distributed amongst different cellular clusters (Suppl. Fig. 4C-H), supporting the versatility of our approach. Importantly, the trans-species data integration relies on a set of genes with uniquely identified homologs in zebrafish and mouse genomes that contain about half of all genes identified in these two species. However, this rudimentary gene set does not compromise the identification of the cellular clusters and their similarities as we identify the same basic cell types containing mouse cells in both integrated and original datasets. Moreover, the set of most variable genes identifying the cell types in the two datasets did not differ significantly. This makes our approach very promising for evolutionary comparisons and we expect it to be even more versatile by comparing more closely related species such as different mammalian species. Although, the basic analysis and the identification of different cellular states are not compromised in our analysis, we cannot exclude that particular cell type specific signaling pathways and regulatory mechanisms are not affected. Therefore, we traced back cells from the integrated data set to the original dataset and used the original gene-set containing all detected genes to address the regulatory pathways in representative populations.

### Molecular features of de-differentiated astrocytes

The de-differentiated astrocytes are the rare population appearing exclusively after a particular type of insult including TBI, bleeding, stroke or epilepsy (Sirko et al., 2013, 2023). Importantly, astrocyte proliferation is the most prominent feature of the plastic astrocytes (Dimou & Gotz, 2014). The gain of plasticity in this set of astrocytes is associated with changes in their cytoarchitecture and up-regulation of intermediate filament GFAP (Escartin et al., 2021; Patani et al., 2023). However, these morphological changes are shared with a number of astrocytic populations that do not gain the proliferation capacity (Sirko et al., 2013). Moreover, a specific manipulation of the innate immunity pathways reduced astrocyte proliferation after brain injury without a change in their morphology or GFAP levels (Koupourtidou et al., 2024). This brings an interesting concept that different aspects of astrocyte reactivity are controlled by different regulatory networks. Our analysis revealed an enrichment of a number of cell specific determinants (*Hmgb2, Uhrf1, Ascl1,* and *Rpa2*) in the de-differentiated astrocytic cluster known for their roles in neural stem cell dynamics, neurogenesis, DNA methylation regulation, and DNA replication/repair (Bostick et al., 2007; Kimura et al., 2018; Păun et al., 2023; Ramesh et al., 2016; Shi et al., 2010; Zhou & Luo, 2013). These molecular features allowed the de-differentiated astrocytes to cluster with zebrafish neural stem cells. However, in stark contrast to zebrafish radial glia (neural stem cells), the de-differentiated astrocytes never give rise to any neurons despite the up-regulation of these neurogenic genes. Our integration now allowed us to directly compare cells from zebrafish and mouse within the same cluster. This analysis revealed a differential enrichment of known neurogenic signaling pathways: the Notch, IL-6 and Wnt playing an important role in controlling neurogenesis in both zebrafish and mouse developing and adult brain (Arredondo et al., 2020; Dray et al., 2021; Kageyama et al., 2009; Storer et al., 2018; Westphal et al., 2022). Indeed, the Wnt pathway activation in radial glia after optic tectum injury, leading to RG proliferation and neurogenesis in adult zebrafish has already been described (Shimizu et al., 2018). These findings are very well in line with the capacity of different ECM components to induce the de-differentiation of astrocytes isolated from the intact brain *in vitro*, supporting a concept that inductive signal in the injured environment is missing in the mouse brain. Moreover, the de-differentiated astrocytes were still enriched in glycolytic processes and processes involved in oxidative stress. Oxidative stress has been associated with the trans-differentiation of astrocytes to neurons (Gascon et al., 2015, 2017). The fate conversion of astrocytes to neurons requires the metabolic switch to oxidative phosphorylation and the mouse de-differentiated astrocytes might fail to do so and as a consequence die. In contrast, the zebrafish stem cells could change their metabolism and generate new neurons in response to injury. This is in line with the transplantation experiments of reactive astrocyte-derived neurospheres into the SEZ that failed to yield neurons (Shimada et al., 2012), suggesting a cell intrinsic block in the lineage.

### A subset of astrocytes incompletely goes through the neurogenic lineage in response to injury

As the comparison of zebrafish and mouse cells from the de-differentiated astrocytic clusters Astro/RG 3/6 suggests an intrinsic barrier for neurogenesis from de-differentiated astrocytes, we integrated these de-differentiated astrocytes to the bona fide neurogenic lineage from the subependymal zone. To our surprise, at least a proportion of de-differentiated cells clustered with TAPs. Importantly, these progenitors have been up-regulating transcription factors such as Olig2 involved in gliogenesis (Nishiyama et al., 2021), suggesting their glial identity. Indeed, such gliogenic TAPs have been previously reported in the neurogenic zone (Colak et al., 2008; Hack et al., 2005; Malatesta et al., 2003; Ortega et al., 2013). These data are in line with our fate mapping experiments using the Ascl1^CreERT2^ mouse line. According to these experiments, the Ascl1-positive de-differentiated astrocytes generate unipotent, gliogenic neurospheres. The analysis of the de-differentiation trajectory of reactive astrocytes along with neurogenic lineage revealed that they go through the activated stem cell-like state in order to generate the TAP-like state. This stem cell like state could then be the possible source of multipotent neurospheres generated from the de-differentiated astrocytes (Buffo et al., 2008; Sirko et al., 2013). Interestingly, the comparison between the stem cell-like astrocytes and bona fide astrocytes revealed an enrichment of inflammatory genes in the stem cell-like astrocytes suggesting that these could be interfering with the neurogenic trajectory. Indeed, the TAP-like cells generated from these inflammatory signature-enriched astrocytes failed to up-regulate the typical neurogenic fate determinants such as Sox4 and Sox11. The upregulation of these factors downstream of the chromatin remodeling factors such as Brg1 is necessary for the completion of the neurogenic cascade and generation of neuroblasts (Ninkovic et al., 2013). Instead, the Brg1-deficient cells generate gliogenic oligodendrocyte progenitors similar to de-differentiated astrocytes. One possibility is that the neurogenic fate is not fully induced or maintained due to increased level of Notch seen in these TAP-like cells of injured cortex (Santopolo et al., 2020; Zamboni et al., 2020). Notch signaling depletion in cortical astrocytes following TBI has been demonstrated to trigger a neurogenic response (Zamboni et al., 2020), possibly linking the intrinsic fate barriers with the inductive signals from the injured environment.

## Methodology

### Source of transcriptome data

We harnessed single-cell transcriptome datasets from our prior investigations, specifically zebrafish data by Zambusi et al., 2022 (GSE179134: Telencephalon, Wt Intact; Telencephalon, Wt 3 dpi; Telencephalon, Wt 7 dpi), mouse data by Koupourtidou et al., 2024 (GSE226207: Intact, bio rep 1; Intact, bio rep 2; 3dpi_CTRL, bio rep 1; 3dpi_CTRL, bio rep 2; 5dpi_CTRL, bio rep 1; 5dpi_CTRL, bio rep 2; 5dpi_CTRL, bio rep 3), mouse adult subependymal zone (SEZ) data from [accession number pending], and RNA-seq data pertaining to astrocyte dedifferentiation from the study conducted by Schmid et al. in 2016 (GSE75589: AC1-RNA; AC2-RNA; TRP1-RNA; TRP2-RNA). Additionally, we incorporated mouse postnatal day 4 cortex data from Di Bella et al. in 2021 (GSE153164: RNA-seq P4). For comparison of integration analysis with different mouse lineages, scRNA-seq data of PBMCs (Peripheral Blood Mononuclear Cells) was sourced from C57BL/6 mice (v1), Single Cell Immune Profiling Dataset by Cell Ranger 3.1.0, 10x Genomics.

### Transcriptome data analysis

Datasets from both mouse and zebrafish under both injured and intact conditions were subjected to initial processing using Seurat package in R. A Seurat object was constructed using the unique molecular identifier (UMI) count matrix with a minimum of 3 cells and 200 genes as cutoffs. In both species’ datasets, cells exceeding 20% mitochondrial reads, featuring RNA counts beyond 6000 or below 200, or having RNA counts less than 40000 were systematically excluded to filter low-quality cells and potential outliers, ensuring the reliability of subsequent analyses. Potential doublets were removed using DoubletFinder (version 2.0.3) package. Normalization and identification of highly variable features were carried out using Seurat’s default parameters. The heterogeneity associated with the cell cycle genes, mitochondria and ribosomal percentage were regressed out using the ScaleData function, taking features as all the genes. Subsequently, a Principal Component Analysis (PCA) was conducted on the resulting matrix. This PCA output was then utilized for Louvain cell clustering and Uniform Manifold Approximation and Projection (UMAP) visualization, providing a comprehensive view of the cellular landscape at 0.6 resolution and dimension 1:15. To identify the differentially expressed genes (DEGs) that serve as cluster biomarkers, we used the FindAllMarkers function of the Seurat package. The DEGs specific clusters between mouse and zebrafish were visualized using function do_VolcanoPlot of SCpubr package. In addition, we scored the known cell-type-specific markers using the Seurat AddModuleScore function and visualized the results using the FeaturePlot function of Seurat and the EnrichHeatmap function of the ScPurb package. The unintegrated mouse and zebrafish species datasets were annotated based on published studies by Zambusi et al. 2022 and Koupourtidou et al. 2024, respectively. Similarly, the mouse SEZ scRNA seq (at resolution 0.8 and dimensions 1:20), postnatal day 4 cortex (at resolution 0.7 and dimensions 1:10) and PBMCs data (at resolution 0.8 and dimensions 1:20), was analyzed and visualized. The RNA seq data of astrocyte dedifferentiation was procured from iDEP 0.96 tool (http://149.165.154.220/idep/) to get log normalized transcript which was further visualized by scaling using pheatmap (version 1.0.12) R package.

### scRNA-seq Integration analysis

We conducted trans-species integration of single-cell RNA sequencing data from both mouse and zebrafish using the Seurat package. Seurat v4 uses canonical correlation analysis (CCA) to identify correlated variables between datasets, with mutual nearest neighbors (MNN) serving as anchor points for integration. Homologous genes between mouse and zebrafish were identified using homologene R packages (version 1.4.68.19.3.27), and a custom ‘gene_convert’ function (https://github.com/NinkovicLab/cross-species-integration) was created to ensure consistent gene nomenclature, taking mouse as a reference. To perform integration, we identified common anchors between the datasets of both species using Seurat’s FindIntegrationAnchors. These anchors were used to integrate the two datasets with the IntegrateData function. Subsequently, the integrated dataset underwent dimensionality reduction PCA, clustering with the Louvain algorithm (at resolution 0.7 and dimensions 1:10), and visualization via UMAP. Differentially expressed features (cluster biomarkers) were identified using the FindAllMarkers function. Similarly, the PBMCs data from mouse integrated using Seurat with Intact and injured cortex of mouse and telencephalon of zebrafish. Furthermore, to enable comparative integrated analysis, we utilized the Harmony package (version 1.1.0) in R, which employs an iterative method for integration, following the guidelines outlined at https://portals.broadinstitute.org/harmony/articles/quickstart.html using similar parameters as Seurat dimension (at resolution 0.7 and dimensions 1:10). The Integration of the cortex and the SEZ regions of mouse was also performed and analyzed in similar way (at resolution 0.8 and dimension 1:30) in order to access the similarities of identified dedifferentiated astrocytic cluster with bonafide neuronal stem/progenitor cells of SEZ. All codes for visualization will be provided upon request.

### Cell distribution plots

To visualize the cell distribution between/within conditions or species or samples, we used various plots like bar plots, alluvial plots, chord diagram plots, and pie charts; generated in Rstudio (version 4.2.3) using ggplot2 (version 3.4.2), DittoSeq using dittoBarPlot function (version 1.8.1) and SCpubr (do_ChordDiagramPlot function) (version 1.1.2) from Seurat object in R. The color palette used in these plots was generated by the Rcolorbrewer (version 1.1-3) package in R.

### Relative similarities heatmap

We assessed the relative similarity between two scRNA-seq datasets using ELeFHAnt (https://github.com/praneet1988/ELeFHAnt) in R. We used the DeduceRelationship function, which predicts the relationship between the datasets based on their gene expression profiles. We used the following default parameters: varfeatures = 2000 (most variable features to use for dimensionality reduction and clustering), classifier = SVM (algorithm to train a classifier on a subset of the data and test it on another subset; shown ∼85% accuracy), and downsample = 200 (randomly samples 200 cells from each dataset to balance the class sizes and reduce the computational cost). The DeduceRelationship function returns a score that indicates how similar the two datasets are, ranging from 0 (no similarity) to 1 (high similarity). These scores can be used to identify cell types that are similar between the two datasets and to compare gene expression patterns across different cell types.

### Pseudotime trajectory and diffusion map analysis

The pseudotime trajectory analysis was performed using monocle3 as described https://cole-trapnell-lab.github.io/monocle3/ in Rstudio. For the analysis we first imported the Seurat object clusters into Monocle3 as cds object. The cells were then ordered along a pseudotime trajectory using the orderCells function, taking homeostatic clusters (in context to integrated astrocytic clusters) and qNSC (in context to integrated SEZ clusters). We visualized the pseudotime trajectory of cells using plot cells function and color pallet by RColorBreweR package. Additionally, we utilized the plot_gene_in_pseudotime function to discern patterns in gene expression along the trajectory for a specific set of genes. To identify the major cell types or states in different conditions, we performed a diffusion map analysis on the Seurat clusters using the “DiffusionMap” function from the destiny package (version 3.1.1) in R with default parameters. We visualized the diffusion map using a scatter plot against the first diffusion component, which captures the main variation of the data. This allowed us to show how cells transition between different states in different conditions, where each point represents a cell and the color indicates the clusters.

### Tracing back cell identity

To trace back the origin of cells from clusters from one object to an integrated object, we extracted the cells of clusters using the WhichCells function of Seurat. These cells were preprocessed using the substring function of R to match the UMI of cells. To visualize the cross-referenced cells, we utilized the highlight.cells function from DimPlot of Seurat. We used DittoBarPlot from Dittoseq to quantify and plot the number of cross-referenced cells with respect to clusters.

### Biological processes and wiki pathway analysis

We used Metascape 3.5 (https://metascape.org), an online tool, to perform biological processes and WIKI pathway analysis on our gene list. We uploaded our gene list using the mouse species and opted for custom analysis, where we specified the following parameters: 1) The annotation was performed using the default databases, including the Gene Ontology (GO) Biological Process and the WIKI Pathway, 2) The enrichment analysis was performed using a hypergeometric test with a p-value cut-off of 0.01, a minimum overlap of 3 genes, and a minimum enrichment of 1.5 for GO biological process and WIKI pathway, and 3) visualization was opted using heatmaps, which showed the expression levels of the genes in each term or pathway across conditions or clusters.

### Animals

All surgeries were performed on 8-15 week old male mice (Mus musculus), housed, and handled under the German and European guidelines for the use of animals for research purposes. The room temperature was maintained within the range of 20–22 °C, while the relative humidity ranged between 45–55%. The light cycle was adjusted to 12 h light:12 h dark period. Room air was exchanged 11 times per hour and filtered with HEPA-systems. All mice were housed in individually ventilated cages (2-5 individuals per cage) under specified-pathogen-free conditions with food (standard chow diet) and water ad libitum. The cages were equipped with nesting material, a red corner house and a rodent play tunnel. Soiled bedding was removed every 7 days. For IHC and in situ experiments, wild-type C57BL/6J animals (strain #000664) were used, while neurospheres assay was performed in the Ascl1^CreERT2^ mouse ((Ascl1^tm1.1(Cre/ERT2)Jejo^; JacksonLab strain 012882) crossed to the tdTomato reporter mouse line ((Ai14; B6.Cg-Gt(ROSA)^26Sortm14(CAG-tdTomato)Hze^; JacksonLab strain 007914)). All animal work was performed in accordance with the German and European Union regulations and approved by the Institutional Animal Care and Use Committee (IACUC) and the Government of Upper Bavaria (AZ: ROB-55.2-2532.Vet_02-20-158). Anesthetized animals received a stab wound lesion in the cerebral cortex by inserting a thin knife (19G, Alcon #8065911901) into the grey matter using the following coordinates from Bregma: RC: –1.2; ML: 1.2 and from Dura: DV: –0.6 mm. To produce stab lesions, the knife was moved over 1mm back and forth along the anteroposterior axis from –1.2 to –2.2 mm as described before. Animals were euthanized 3 and 5 days after the injury (dpi) by transcardial perfusion (for more details see section tissue preparation). For the induction of Cre-mediated recombination in Ascl1^CreERT2^ crossed to tdTomato reporter mice, tamoxifen (40 mg/ml, Sigma #T5648) was administered orally (20G, Merck #CAD9921). Animals received tamoxifen twice (400 mg/kg per treatment).

### BrdU labeling

Proliferating cells were labeled *in vivo* via water administration of the thymidine analog 5-bromo-2’-deocyuridine (BrdU). To this end, BrdU (1 mg/mL) and sucrose (1 %) were added to the animals’ drinking water starting from 24h after injury until euthanasia. Water was exchanged every 2 days.

### Tissue preparation

Mice were deeply anesthetized and transcardially perfused with phosphate-buffered saline (PBS) followed by 4% paraformaldehyde (PFA) (wt/vol) dissolved in PBS. Brains were postfixed in 4% PFA overnight at 4 °C, washed with PBS and cryoprotected in 30% sucrose (Carl Roth #4621.2) at 4 °C for IHC. For RNAscope® in situ hybridization (ISH), brains were incubated in gradually concentrated sucrose solutions at 4 °C, starting with 10 % sucrose in 1X PBS, followed by 20 % and finally 30 % sucrose in 1X PBS. Brains were embedded in frozen section medium Neg-50 (Epredia #6502), frozen and subsequently sectioned using a cryostat (Thermo Scientific CryoStar NX50). Coronal sections were collected either at a thickness of 20 μm on slides for RNAscope (Epredia #J1800AMNZ) or 40 μm for free-floating immunohistochemistry.

### Immunohistochemistry

For immunohistochemistry, sections were blocked and permeabilized with 10% normal goat serum (NGS, vol/vol, Biozol #S-1000)/donkey serum (NDS, vol/vol, Sigma-Aldrich #566460) and 0.5% Triton X-100 (vol/vol, Sigma-Aldrich #T9284) dissolved in 1xPBS. The same solution was used to dilute the primary antibodies. Primary antibodies were incubated with sections overnight at 4 °C. Following primary antibodies were used: anti-RPA32/RPA2 (rabbit IgG, 1:250, Abcam, ab76420), anti-HMGB2 (rabbit IgG, 1:1000, Abcam, ab67282), anti-RFP (rabbit IgG, 1:1000, Rockland/Biomol, 600-401-379), anti-BrdU (rat IgG2a, 1:500, Abcam, ab6326), anti-GFAP (for IHC: mouse IgG1, 1:500, Sigma-Aldrich, G3893 and for combination with RNAScope: goat IgG, 1:250, Abcam, ab53554); anti-DCX (guinea pig, 1:1000, Merck/Millipore, AB2253), anti-O4 (mouse IgM, 1:50, Sigma, O7139). Sections were washed with PBS and incubated with secondary antibodies dissolved in blocking solution (10% NGS/ NDS and 0.5% Triton-X-100 in PBS) for 2 h at room temperature. Following secondary antibodies were used: goat anti-mouse IgG1 Alexa Fluor™ 488 (1:1000, Thermo Fisher Scientific A21121), goat anti-rabbit IgG Alexa Fluor™ 546 (1:1000, Thermo Fisher Scientific A11035), goat anti-rat IgG Alexa Fluor™ 647 (1:1000, Thermo Fisher Scientific A21247), donkey anti-goat IgG Alexa Fluor™ 488 (1:1000, Thermo Fisher Scientific A11055), donkey anti-rabbit IgG Alexa Fluor™ 594 (1:1000, Thermo Fisher Scientific A21207), donkey anti-Rabbit IgG Alexa Fluor™ 647 (1:1000, Thermo Fisher Scientific A31573), goat anti-mouse IgG2a A488 (1:1000, Thermofisher A21131). For nuclear labelling, sections were incubated with DAPI (final concentration of 4 µg/mL, Sigma #D9542) for 10 min at room temperature. Stained sections were mounted on glass slides (Epredia #AG00000112E01MNZ10) with Aqua-Poly/Mount (Polysciences #18606). For BrdU detection, sections were pre-treated with HCl (4 N), followed by three washes using borate buffer (0.1 M) and another three washes with 1X PBS before incubation with primary antibody solution. For RPA2 staining, antigen retrieval using Dako TRS (Agilent, Dako S1699) was performed prior to primary antibody incubation. Dako solution was first diluted 1:10 in distilled water (diH_2_O) and then prewarmed at 65 °C for 15-20 minutes. Sections were incubated in the diluted DAKO solution at 95 °C for 20 minutes followed by another 15 minutes at 65 °C to slowly cool down. After cooling-down to room temperature, sections were washed three times in 1X PBS and incubated in primary antibody solution.

### In situ hybridization

RNA in situ hybridization was performed using RNAscope® Multiplex Fluorescent Reagent Kit (ACD, 323110) according to the manufacturer’s instructions. Briefly, brain sections were airdried, shortly washed in 1X PBS to remove remaining OCT, baked at 60°C for 30 minutes and then post-fixed in 4 % PFA at 4°C for 15 minutes. Next, sections were ethanol-dehydrated (Carl Roth #9065.4), treated with H_2_O_2_ (ACD, 322381) and protease-permeabilized for 20 min at 40 °C. Brain sections were then incubated for 2 h at 40 °C using the following probes: RNAscope® Probe –Mm-Uhrf1 (Bio-Techne 559891) and RNAscope® Probe –Mm-Ascl1-CDS-C3 (Bio-Techne 476321-C3). The signal was amplified using Cyanine 3 Plus Amplification Reagent and the Cyanine 5 Plus Amplification Reagent from Akoya Biosciences (NEL744001KT and NEL745001KT) always diluted 1:750 in 1X Plus Amplification Diluent (according to the manufacturer’s instructions). Following washings steps with 1xPBS, sections were fixed for 15 min in 4% PFA at 4 °C and subjected to immunohistochemistry analysis as described above.

### Neurosphere assay

Neurosphere cultures were prepared as previously described (Buffo et al., 2008) using a volume of tissue punched (B0.35 cm) from the lesioned areas of the somatosensory cerebral cortex obtained from the injured brains 5 days after injury. After removal of meninges and white matter, grey matter cells were plated at a density of one cell/ 10 microliters (clonal density) in 500 microliters of neurosphere medium with FGF2 and EGF (both at 20 ng/ml, Invitrogen). The number of neurospheres and the expression of the reporter was assessed after 14 days. The individual neurospheres were assessed for their differentiation capacity by plating individual neurospheres on the PDL-coated coverslips as described previously (Buffo et al., 2008).

### Image acquisition and processing

Confocal microscopy was performed at the core facility bioimaging of the Biomedical Center (BMC) with an inverted Leica SP8 microscope using the LASX software (Leica). Overview images were acquired with a 20x/0.75 objective, higher magnification pictures with a 40x/1.30 or 63x/1.40 objective, respectively. Image processing was performed using the NIH ImageJ software (version 2.9.0/1.53t). For quantifications, a minimum of three sections per animal was analyzed for five animals in total. In each section, an area of 300 μm was selected around the injury (150 μm on either side) and the number of positive cells in all individual z-planes of the optical stack was quantified using the Fiji plug-in tool ‘Cell Counter’.

## Supplementary Figures legends

**Supplementary Figure S1.**
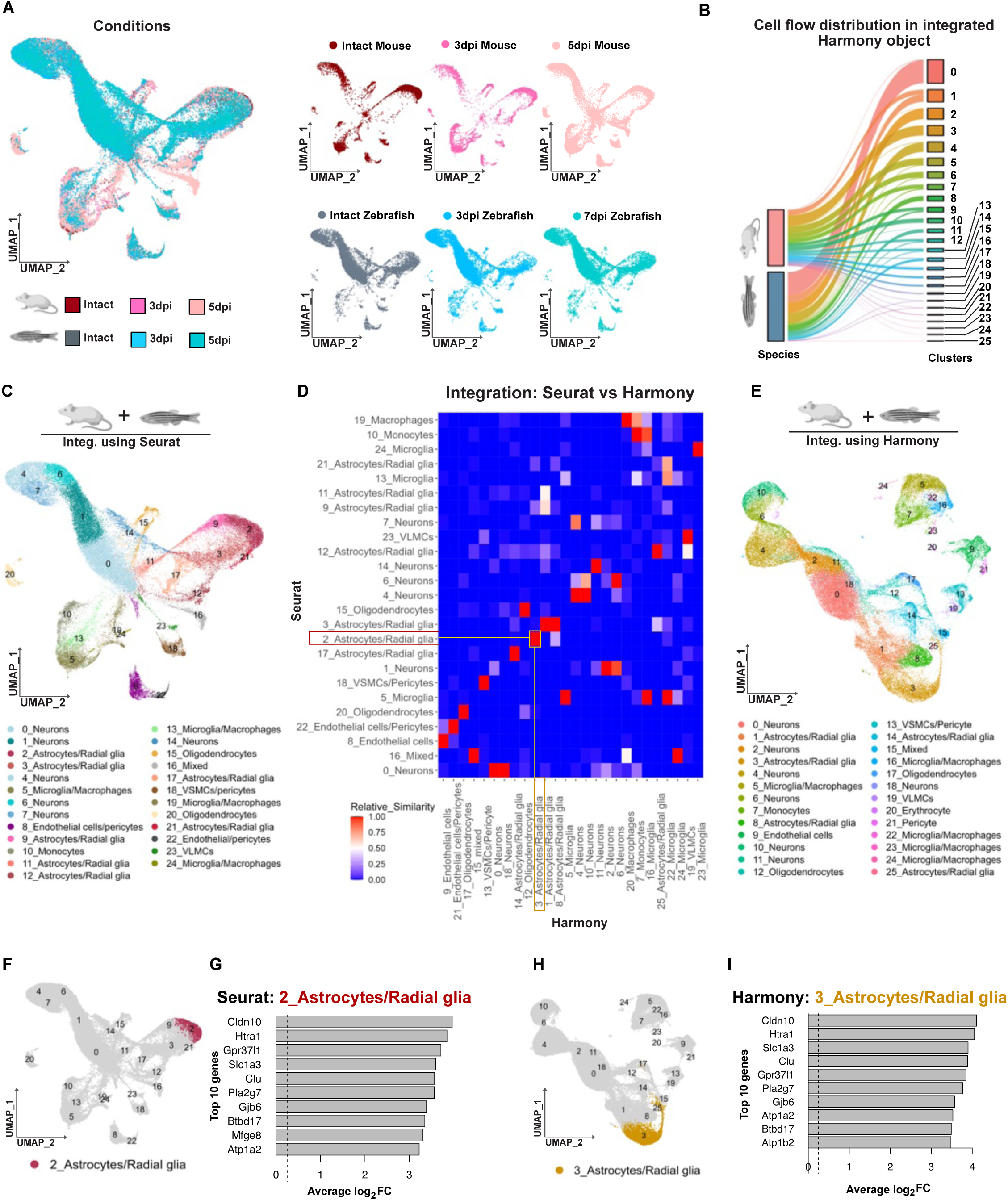
Comparison of Harmony– and Seurat-based integration of mouse and zebrafish datasets. (**A**) UMAP plots depicting Seurat-based integration of cells from injured and intact mouse and zebrafish brain. (**B**) Alluvial plot showing the distribution of mouse and zebrafish cells amongst different clusters following harmony integration. **(C, E)** UMAPs depicting cellular clusters with their identity after Seurat (C) and harmony (E) based integration. (**D**) Relative similarity heatmap comparing integrated and annotated clusters by Seurat and harmony. Highlighted 3_Astrocytes/Radial cluster (yellow) in harmony analysis corresponds to 2_Astrocytes/Radial cluster (red) from Seurat integration. (**F**, **H**) UMAP plots highlighting clusters 2_Astrocytes/Radial cluster in the Seurat integration (F) and corresponding cluster 3_Astrocytes/Radial cluster in Harmony based integration (H). (**G**, **I**) Bar plots depicting the enrichment (log_2_FC) of top 10 enriched genes in the corresponding 2_Astrocytes/Radial cluster in Seurat analysis (G) and 3_Astrocytes/Radial cluster in harmony analysis (I).

**Supplementary Figure S2.**
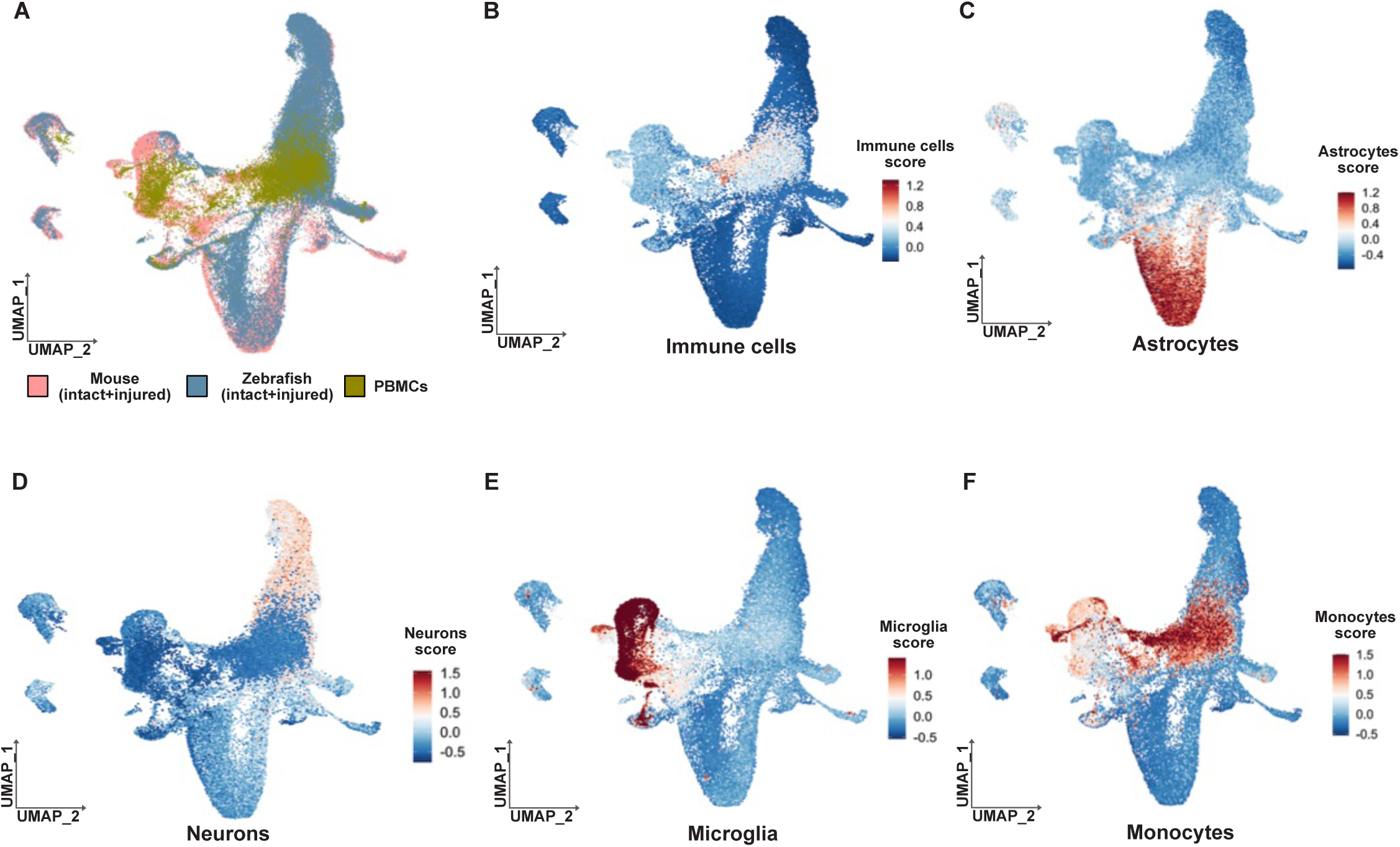
Integration of brain datasets with the dataset of peripheral blood mononuclear cells (PBMCs). (**A**) UMAP plot depicting integration of mouse (intact+3dpi+5dpi), zebrafish (intact+7dpi+7dpi) and PBMCs cells. (**B**-**F**) UMAP plots showing expression score for immune cells (B), astrocytes (C), neurons (D), microglia (E) and monocytes (F) in integrated dataset. Gene lists used for the expression score generation are provided in the Suppl. Table 1 and Suppl. Table 2.

**Supplementary Figure S3.**
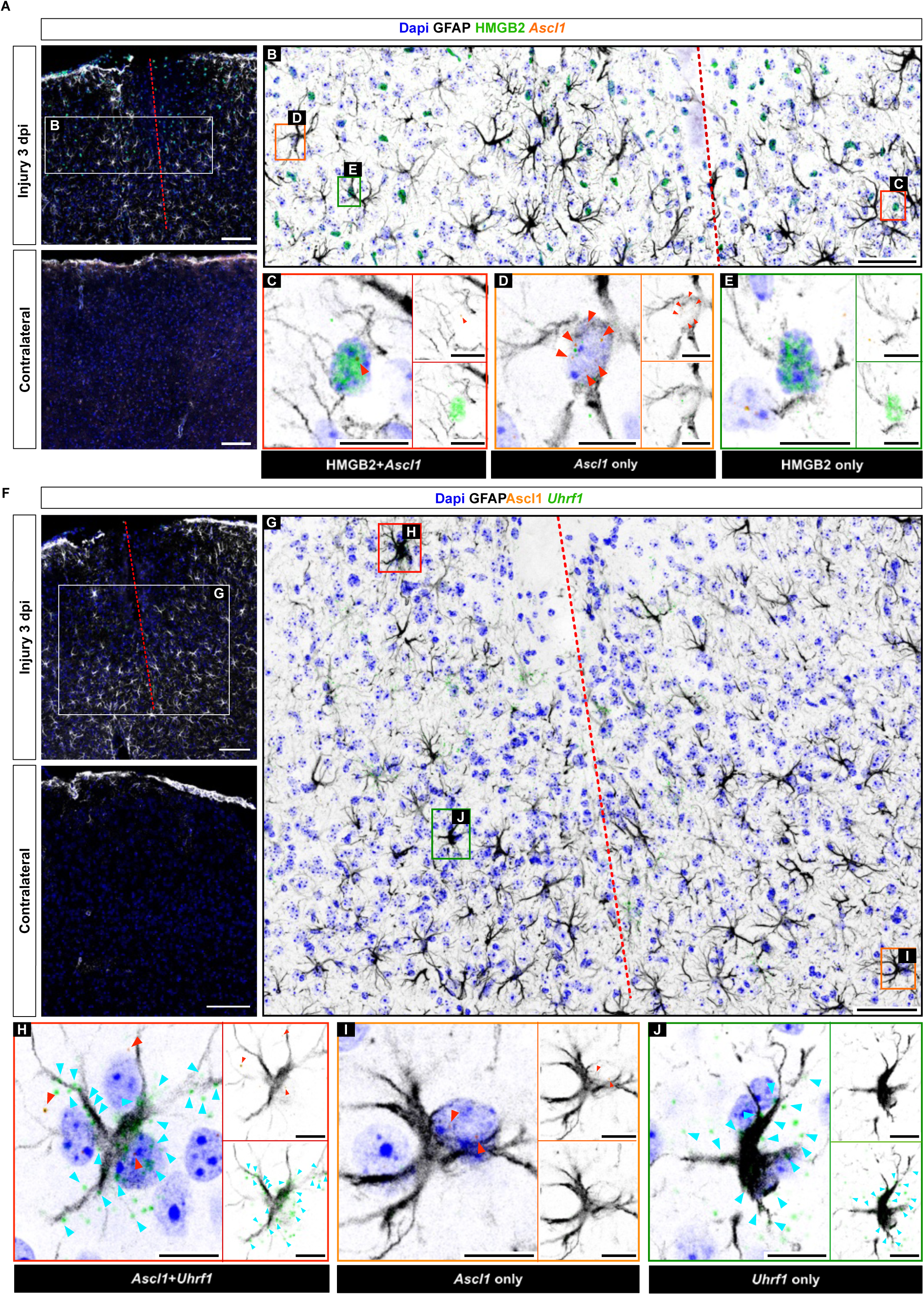
Expression of Astro/RG_6/3 enriched genes in reactive astrocyte population. (**A**, **F**) Micrographs depicting GFAP, HMGB and *Ascl1* expression in the injured (upper panel) and intact (lower panel) cerebral cortex at 3 dpi. (**B**-**E**) Micrographs showing HMGB2 expression and the RNAscope^®^ signal for *Ascl1* in reactive, GFAP+ astrocytes 3 dpi. (**F**, **J**) Micrographs illustrating the RNAscope^®^ signal for *Ascl1* and *Uhrf1* in reactive, GFAP+ astrocytes 3 dpi. Micrographs in B and G are magnifications of areas boxed in A and F. Micrographs in C-E and H-J are magnifications of cells boxed in B and G according to the color code. Red arrowheads in single plane views (C-E and H-J) indicate Ascl1 molecules and cyan arrowheads point at *Uhrf1* molecules visible in the cell within that visual plane. Dashed lines indicate injury site. Micrographs A, B, F and G are maximum intensity projections of the confocal Z-stack. Micrographs C-E and H-J are single optical sections. Scale bars in A, F 100 μm; in B, G 50 μm; in C-E and H-J 10 μm.

**Supplementary Figure S4.**
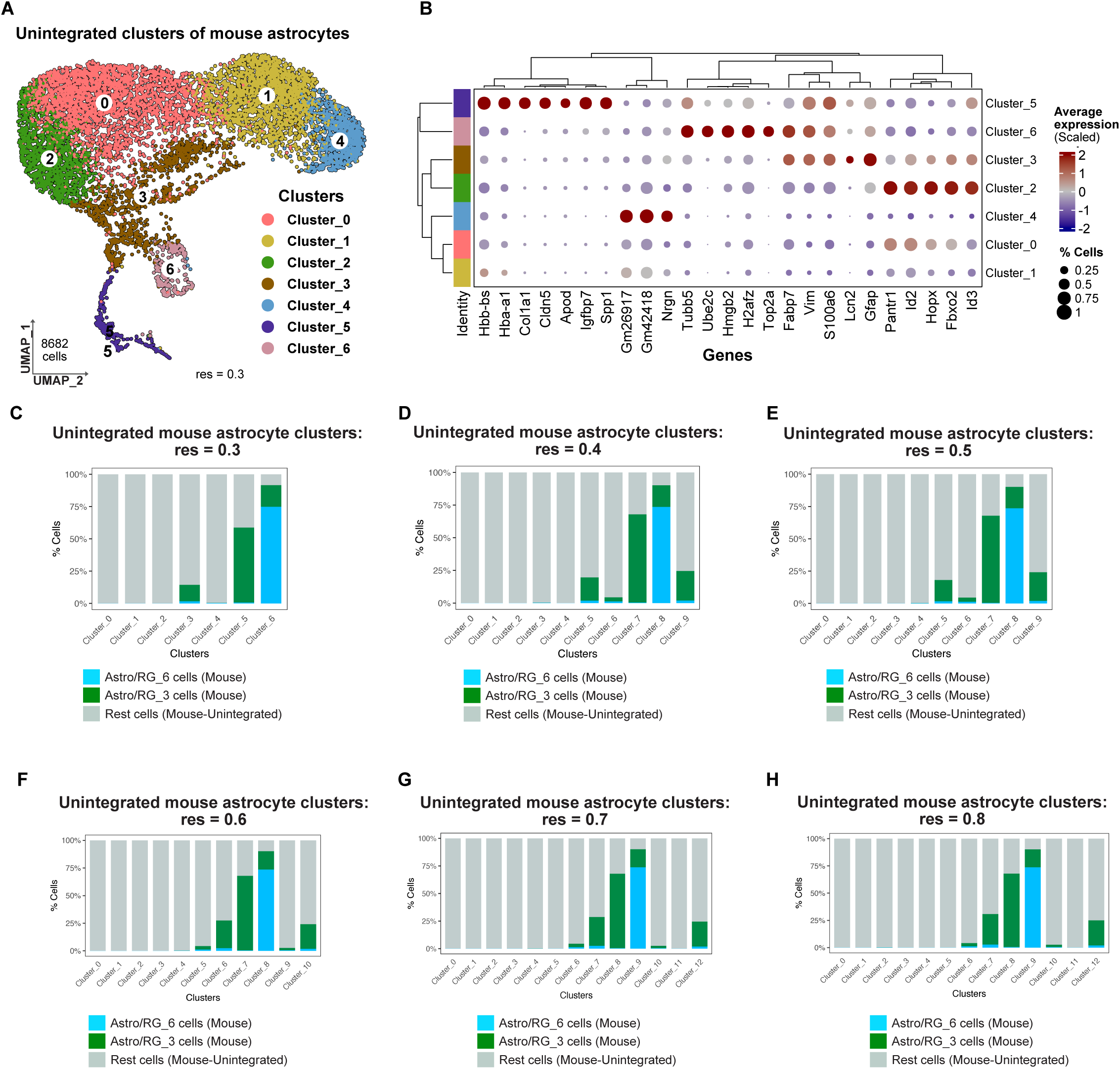
Cells from the Astro/RG 3 and 6 clusters are dispersed amongst different astrocytic clusters in unintegrated mouse dataset. (**A**) UMAP plot depicting 7 distinct astrocytic clusters at resolution 0.3. (**B**) Dot plot highlighting the top 5 expressed genes in each astrocytic cluster shown in A, color-coded by expression levels. (**C**-**H**) Bar plots highlighting distribution of Astro/RG clusters 3 (green) and 6 (blue) cells across astrocytes clusters in unintegrated dataset at resolutions of 0.3 (C), 0.4 (D), 0.5 (E), 0.6 (F), 0.7 (G), and 0.8 (H).

**Supplementary Figure S5.**
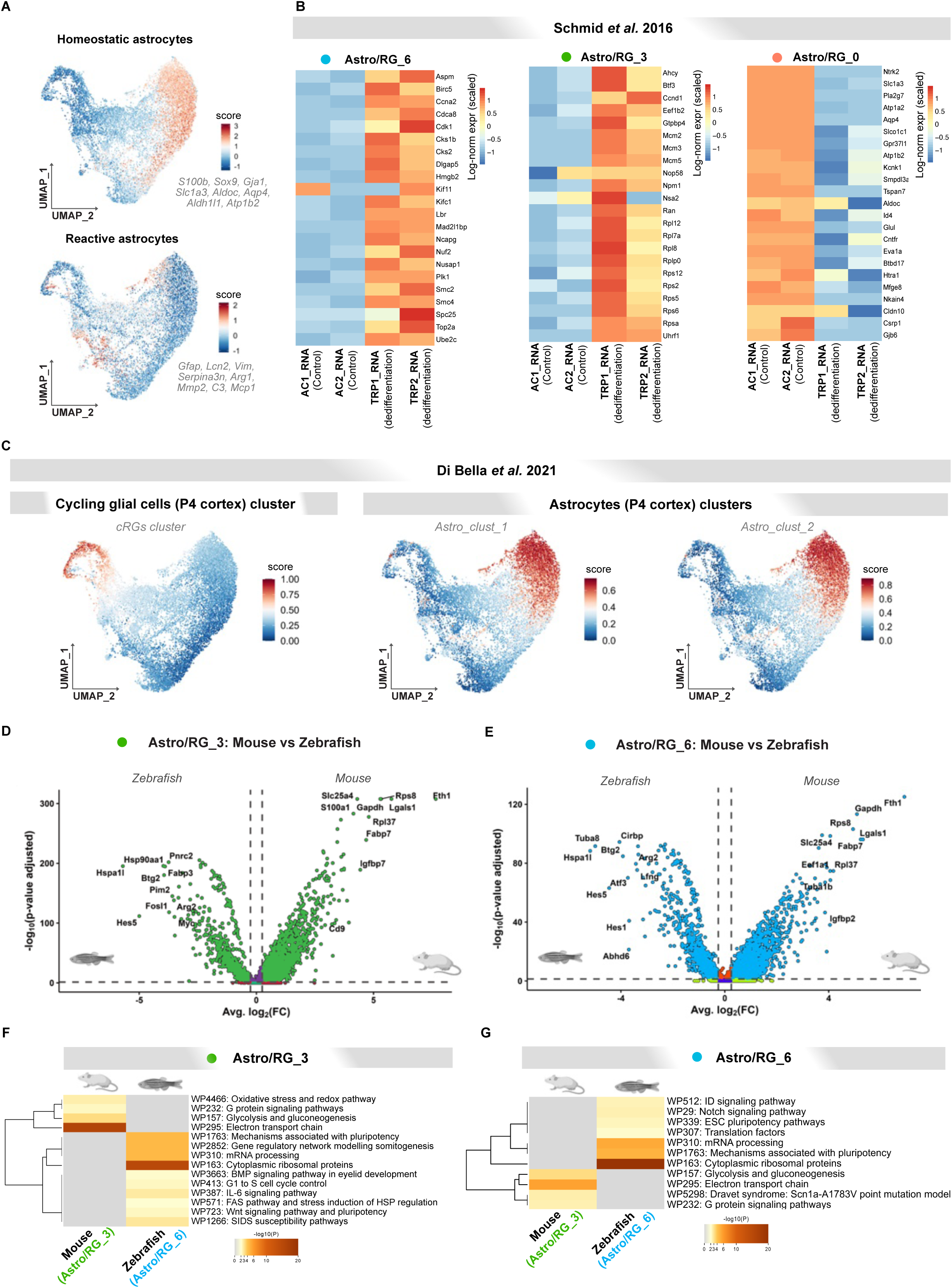
Injury-induced mouse Astro/RG 3 and 6 cluster cells share molecular features with immature astrocytic progenitors. (**A**) UMAP plots depicting the expression scores according to genes enriched in homeostatic and reactive astrocytes (based on the classification in Koupourtidou et al. 2024) in integrated astrocytic clusters. (**B**) Heatmaps depicting expression of genes identifying injury-induced (Astro/RG 6 and Astro/RG 3) clusters and homeostatic (Astro/RG 0) cluster in control (AC samples) and dedifferentiated (TRP samples) astrocytes. The astrocyte data are obtained from the Schmid et al. dataset (Schmid et al., 2016). (**C**) UMAP plots illustrating the gene expression scores identifying cycling radial glia and astrocytes isolated from postnatal day 4 (P4) mouse cortex in integrated mouse and zebrafish Astro/RG clusters. The P4 dataset comes from Di Bella *et al*. 2021. (**D**, **E**) Volcano plots depicting DEGs between mouse and zebrafish cells in Astro/RG 3 (D) and Astro/RG 6 (E) cluster. (**F**, **G**) Heatmaps depicting enriched GO terms in the set of DEGs between mouse and zebrafish cells in Astro/RG 3 cluster (F) and Astro/RG 3 cluster (G), color-coded by p-values.

**Supplementary Figure S6.**
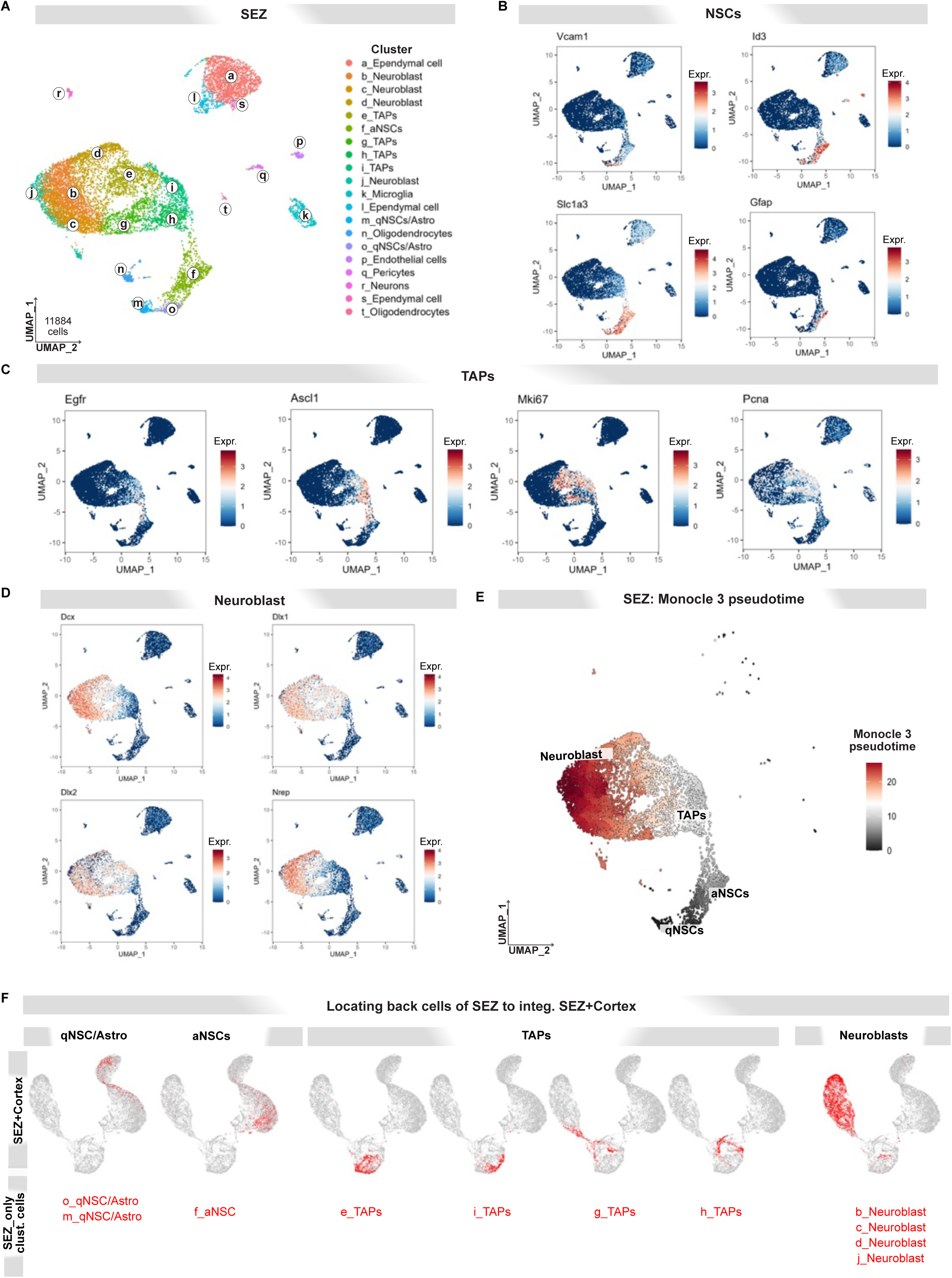
Identification of SEZ cell types and their differentiation trajectories. (**A**) UMAP plot depicting SEZ cells grouped into 20 distinct transcriptional clusters, annotated by cell type-specific markers (Suppl. Table 1 and 7). (**B**-**D**) UMAP plots depicting the expression of known marker genes used for annotation of NSCs (qNSCs/Astro and aNSCs) (B), TAPs (C), and Neuroblasts (D). (**E**) UMAP plot of pseudotime trajectory starting with qNSC, transiting via aNSC and TAPs, and ending in Neuroblasts clusters of SEZ. (**F**) UMAP plots locating the cells of the SEZ lineage within the integrated SEZ+cortex dataset. Cells from the specific SEZ cluster are marked in red.

**Supplementary Figure S7.**
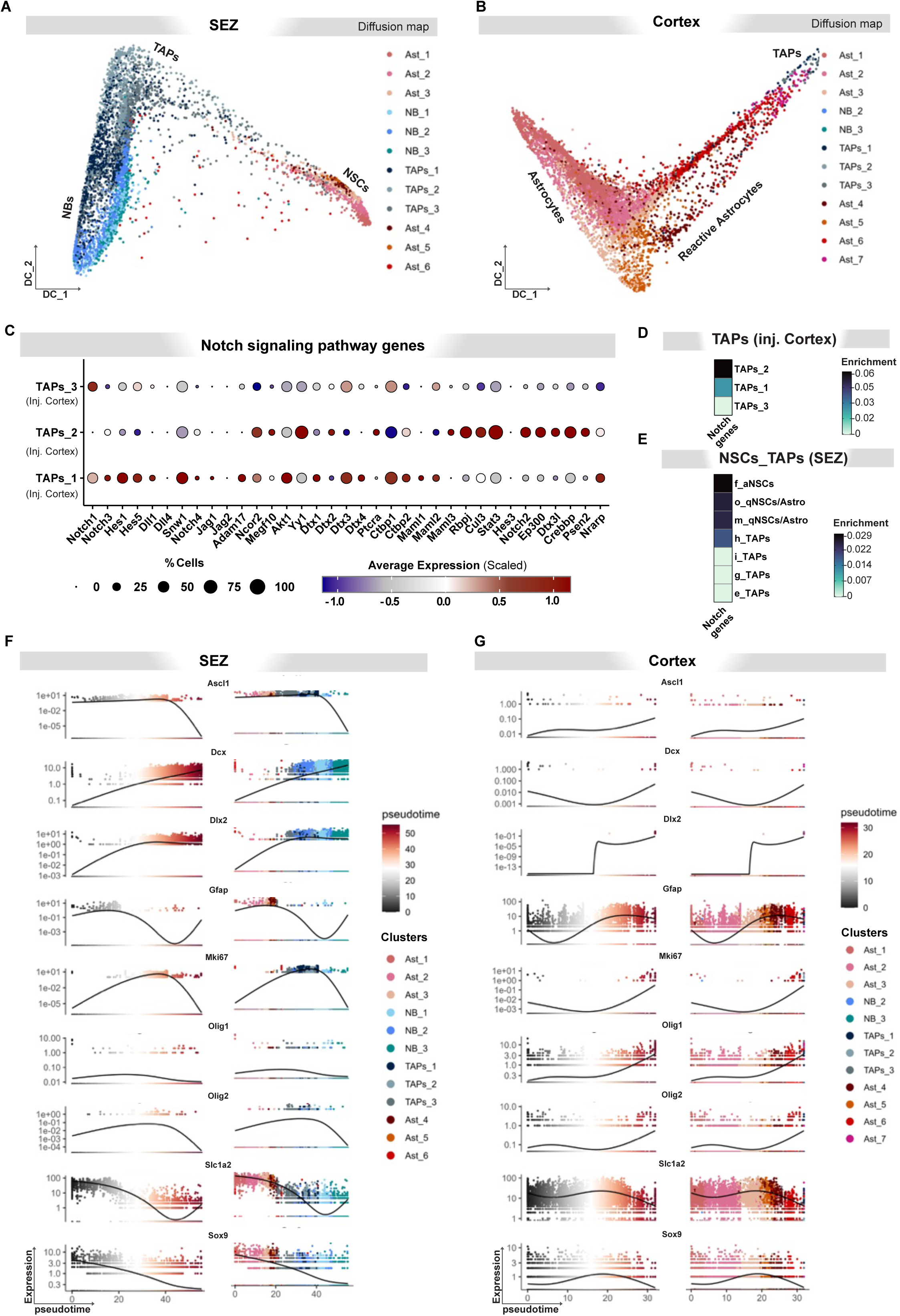
Diffusion map-based definition of the lineage trajectory and gene expression changes along the trajectory. (**A**, **B**) Diffusion component plot of SEZ (A), and cortex (B) from integrated SEZ+cortex dataset displaying the position of Ast, TAPs and NB states. (**C**) Dot plot showing expression of genes downstream of Notch receptor in TAPs clusters from injured cortex. (**D**-**E**) Heatmap plots depicting enrichment expression score for genes downstream of Notch signaling (shown in C) in TAPs clusters from integrated injured cortex (D), NSCs and TAPs clusters from SEZ only dataset (E). (**F**-**G**) Expression dynamics of selected genes along pseudotime trajectory in SEZ (F), and cortex (G) within the integrated SEZ+cortex dataset.

## Supplementary Tables

**Supplementary Tables 1:** Cell type-specific marker genes used for annotation of integrated mouse and zebrafish clusters.

**Supplementary Table 2:** Known immune cell markers used for annotation of peripheral blood mononuclear cells (PBMCs).

**Supplementary Table 3:** This table contains the gene list from the Astro/RG_0 cluster, which includes genes identified as part of the homeostatic clusters of astrocytes (Koupourtidou et al., 2024).

**Supplementary Table 4:** Gene list used to generate score for positive regulation of cell cycle (GO:0045787).

**Supplementary Table 5:** Gene list used to generate scores for cycling glial progenitors and astrocytes clusters of P4 cortex(Di Bella et al., 2021). This includes Astro_clust_1, Astro_clust_2, and Cycling GCs clusters from reanalyzed P4 cortex.

**Supplementary Table 6:** Differentially expressed genes between Astro/RG cluster 3 and 6 cells originating from mouse and zebrafish.

**Supplementary Table 7**: Marker genes used to identify NSC/Astro, TAPs and NBs.

**Supplementary Table**: Differentially expressed genes between TAPs_1 cortical cluster and TAPs_3 SEZ cluster.

**Supplementary Table 8:** Genes involved in the Notch signaling pathway for Mus musculus (house mouse), as identified in the KEGG database.

## Authors contribution

P.M, J.N., M.G. and F.B. conceived the project and designed bioinformatic experiments. C.K., A.Z., J.F.S., K.T.N. generated single cell sequencing data. P.M. performed the bioinformatic analyses. F.B. performed Immunohistochemical and RNAScope analyses. J.N, K.T.N, C.O. S.J, and T.S. performed neurosphere assay. P. M. and J.N. wrote the manuscript with input from all authors.

